# Spatial arrangement and biofertilizers enhance the performance of legume – millet intercropping system in rainfed areas of southern India

**DOI:** 10.1101/2020.11.20.387886

**Authors:** Devesh Singh, Natarajan Mathimaran, Jegan Sekar, Prabavathy Vaiyapuri Ramalingam, Yuvaraj Perisamy, Kathiravan Raju, Rengalakshmi Raj, Israel Oliver King, Thimmegowda Matadadoddi Nanjundegowda, Manjunatha Baiyapalli Narayanswamy, Bhavitha Nayakanahalli Chikkegowda, Savitha Matakere Siddegowda, Davis Joseph Bagyaraj, Paul Mäder, Thomas Boller, Ansgar Kahmen

## Abstract

Biofertilization via the inoculation with arbuscular mycorrhizal fungi (AMF), combined with rhizobia and plant growth promoting rhizobacteria (PGPR), are beginning to become established as an effective and sustainable measure to improve yields. Biofertilization might have a particular potential to boost the yield of intercropping systems in rainfed areas because AMF can form a common mycorrhizal network (CMN) that can transfer nutrients and water between two plants and balance as such belowground competition. In this study, we tested if biofertilizers can enhance the yield of intercropping systems using a pigeon pea (PP) – finger millet (FM) intercropping system grown for two consecutive growing seasons (2016/17 and 2017/18) at two contrasting sites in Bengaluru and Kolli Hills, India. To validate the process of bioirrigation (transfer of water between rhizosphere of two plants), we tested, for the first time, if the spatial arrangement of intercropped plants using either a row-wise or a mosaic design affected yield and water relations with and without biofertilizers. Our results demonstrate that intercropping can improve the straw and grain yield in PP–FM intercropping compared to the respective monocultures but that intercropping effects vary depending on the site characteristic such as climate and soil type. Spatial arrangement of component plants affected the total, straw and grain biomass in intercropping treatments, but this effect also varied across sites. Most importantly, the results from the 2017-18 growing season clearly demonstrated a positive effect of biofertilizer on biomass yield, and this effect was irrespective of site, spatial arrangement, mixed or monoculture. Despite a yield increase in intercropping, we did not see a positive effect of biofertilization on water relations of FM possibly due to interspecific competition for soil moisture where PP dominated. In summary, our study shows the potential of biofertilizers to increase the yield of intercropping systems in rainfed dryland agriculture.

## Introduction

Intercropping has been considered a sustainable way to utilize and share natural resources among different crop species and to improve and stabilize crop yield (Brooker et al., 2015; Martin-Guay et al., 2018). In intercropping systems two or more crop species are grown together (Vandermeer, 1989). Crop yield in intercropping systems are often higher than in sole cropping systems because resources such as soil moisture and nutrients are utilized more efficiently (Lithourgidis *et al*., 2007; Dahmardeh *et al*., 2009; Martin-Guay *et al*., 2018). This is because interspecific competition between intercropping partners is often lower than the intraspecific competition so that a yield advantage occurs (Davis and Woolley, 1993). In addition, beneficial effects of intercropping can come from resource facilitation. As an example, legume–cereal intercropping systems have been widely used in areas with poor soil quality (L Li *et al*., 2007), where legumes fix nitrogen (N) and solubilize phosphorus (P), which is then used by both intercropping partners (Hinsinger *et al*., 2011). In return, cereals can support legumes in two ways, by preventing nitrate-N accumulation in soil which inhibits N fixation by legumes, and by increasing iron availability which enhances N fixation (Zuo *et al*., 2004; Schipanski and Drinkwater, 2012).

In rainfed areas of the arid and semiarid tropics, intercropping has also been suggested to enhance the water availability of shallow-rooted crops via the facilitation of water by deep-rooted plants through hydraulic lift (HL) (Xu, Li and Shan, 2008; Mao *et al*., 2012). The water released from deep-rooted plants due to HL into topsoil layer becomes available to neighbouring shallow-rooted plants, a process termed bioirrigation (Burgess, 2011). The functionality of bioirrigation in intercropping systems has only been tested in a few studies – mainly under controlled conditions in the greenhouse. Sekiya and Yano (2002) showed in a field experiment that pigeon pea (a deep-rooted legume) has the potential to perform HL and could supply deep water to shallow-rooted maize. In another study, Sekiya et al. (2011) showed that plants with deep roots are ideal for intercropping with shallow-rooted crops in water limited agriculture fields and that this kind of intercropping system allows shallow-rooted plant to access deep soil moisture without having deep roots. Other studies have also shown the transfer of hydraulically lifted water (HLW) from a deep-rooted plant to neighbouring shallow-rooted plants (Caldwell and Richards, 1989; Moreira *et al*., 2003; Brooks *et al*., 2006; Bogie *et al*., 2018). While these experiments have suggested that bioirrigation could be an important mechanism for drought stress avoidance of intercropped field crops, evidence for the efficiency of this mechanism in the field is yet lacking.

The success of an intercropping system in the field depends on the avoidance of competitive growth inhibition among the intercropping partners. This requires appropriate spacing of the intercropping partners so that competitive, complementary and facilitative interactions are well balanced and that yield improvements can be achieved. In particular, for bioirrigation to be effective, it seems that an ideal spacing between the intercropping partners is essential. On the one side, intercropping partners have to be arranged with sufficient space among each other in order to avoid competition. On the other side, plants need to be spaced in close enough distance to allow rhizosphere to rhizosphere transfer of bioirrigated water (Burgess, 2011; Prieto *et al*., 2011).

In addition to intercropping approaches, “biofertilization” such as inoculation with arbuscular mycorrhizal fungi (AMF), combined with rhizobia and plant growth promoting rhizobacteria (PGPR), are beginning to become established as an effective and sustainable measure to improve yields (Schütz et al., 2018; Mathimaran et al., 2020, Mäder et al., 2011). The role of AMF for the uptake and transfer of nutrients and water to host plants has been well demonstrated (Augé *et al*., 2001; Querejeta, Egerton-Warburton and Allen, 2003). Biofertilization might have a particular potential to boost the yield of intercropping systems because AMF can form a common mycorrhizal network (CMN) that can transfer nutrients between two plants and balance as such belowground competition (Smith and Read, 2008). In addition, a CMN between the roots of two plants can also constitute a pathway for the transfer of water. Egerton-Warburton et al. (2007) have demonstrated that arbuscular mycorrhizal hyphae provide indeed a potential pathway for the transfer of HLW between two plants. Our recent work has shown that a CMN plays a key role in facilitating the transfer of water between the rhizospheres of two intercropped partners in a greenhouse and can in turn improve the water relations of shallow rooted crops during soil drying (Singh *et al*., 2019). However, a further experiment with bigger pots (50 L) than in the previous experiment did not show an effect of the CMN on water-relations but treatments with CMN had lower foliar damage than treatments without CMN during drought (Singh *et al*., 2020).

The effects of biofertilizers on stabilizing and improving the yields in intercropping systems by improving water relations via bioirrigation have not yet been tested under field conditions, though recent greenhouse studies have shown evidence of facilitation of bioirrigation by AMF and PGPRs (Saharan *et al*., 2018; Singh *et al*., 2019). Furthermore, it is unclear to what extent beneficial effects of biofertilizers in intercropping systems depend on an appropriate spacing of the crops and if – given the appropriate spatial arrangement of crops – the establishment of a CMN can indeed facilitate bioirrigation and improve as such the water relations of shallow-rooted crops in intercropping systems in dryland agriculture. In this study, we investigated the effects of biofertilization on the yield of a legume – cereal intercropping system, and tested different spatial arrangements of the plants in combination with biofertilizer treatments. We used pigeon pea (*Cajanus cajan*) (PP) as a deep-rooted plant and finger millet (*Eleusine coracana*) (FM) as shallow-rooted plant to investigate the following research questions: (i) Does the spatial arrangement of intercropping partners (Fig. 1a) affect straw and grain yield in a FM – PP intercropping system compared to monocultures of the same crops? (ii) Does the application of biofertilizers have an influence on the intercropping effect in spatially differently arranged intercropping systems? (iii) Can intercropping in conjunction with a CMN lead to an improvement of the water relations of shallow-rooted crops?

**Fig. 1a.**
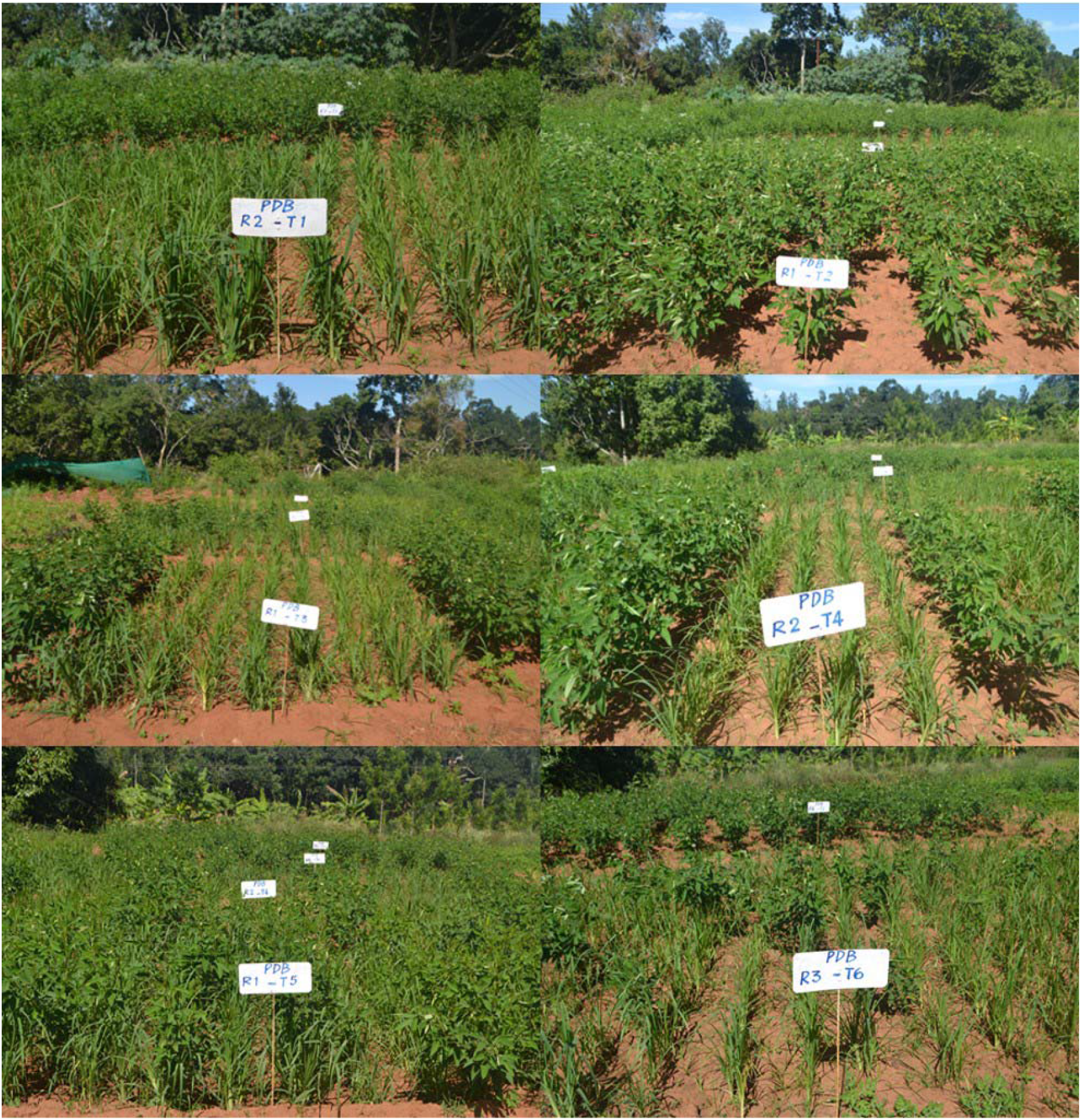
Field trial site at Kolli Hills, India, with various spatial arrangements of component crops pigeon pea (PP) and finger millet (FM) in intercropping systems

## Material and methods

### Selection of field experiment site and crop varieties

To test the influence of the spatial arrangement and biofertilizers on crop yields of PP and FM, field trials were carried out at two different locations during the growing seasons 2016/17 and 2017/18. One experimental site was located at the research field of the University of Agricultural Sciences, Gandhi Krishi Vigyana Kendra Campus (GKVK), Bengaluru, Karnataka. The other site was located at the research field of M S Swaminathan Research Foundation (MSSRF), Kolli Hills, Tamil Nadu, India. Both experimental sites were selected because farmers have already adopted a cereal-legume intercropping system there and have been cultivating PP and FM as one of their main crops. Based on farmers practice in the region and recommendations from local agronomists, we selected FM (GPU-28) and PP (BRG-2) for the field experiment in Bengaluru site while at the Kolli Hills site, PP (SA-1) and FM (Suruttai kelvaragu) were selected.

### Rainfall

The total annual precipitation at Bengaluru site was 694.9 mm in 2016 and 1104.5 mm in the year 2017. At Kolli Hills, the total annual precipitation was 281.7 mm in 2016 and 1690 mm in 2017. Rainfall data recorded during the experimental period indicate that the Kolli Hills area received less rain than Bengaluru site (Fig. S1, supplementary material). Both sites received the maximum amount of rain during the months of May, June and July. Bengaluru site received up to 40-60 mm rain during September, October and December, while Kolli Hills site was completely dry after July during 2016. During year 2016, both research sites, received significantly low precipitation and during few months our weather station at field site recorded very low data. Therefore, to clearly visualize and compare the precipitation, precipitation data from nearest sites as recorded by the Climate Research Unit (Harris *et al*., 2020) have been shown in fig. S1.

### Intercrop field design with different spatial arrangement of PP and FM

The plot size for a treatment was 7.2 x 3.6 m (width x length) with a net plot area of 3.6 x 1.8 m (Fig. 1b). The net plot area defines the central part of each plot as marked in fig. 1b, where all physiological, growth and yield parameters were assessed. The field experiments had six treatments: FM monoculture (T1), PP monoculture (T2), 2:8 (PP:FM) row-wise intercropping (T3), 1:4 (PP:FM) row-wise intercropping (T4), 100% mosaic (T5) and 50% mosaic (T6) (Fig. 1b). Each treatment was replicated four times.

**Fig. 1b.**
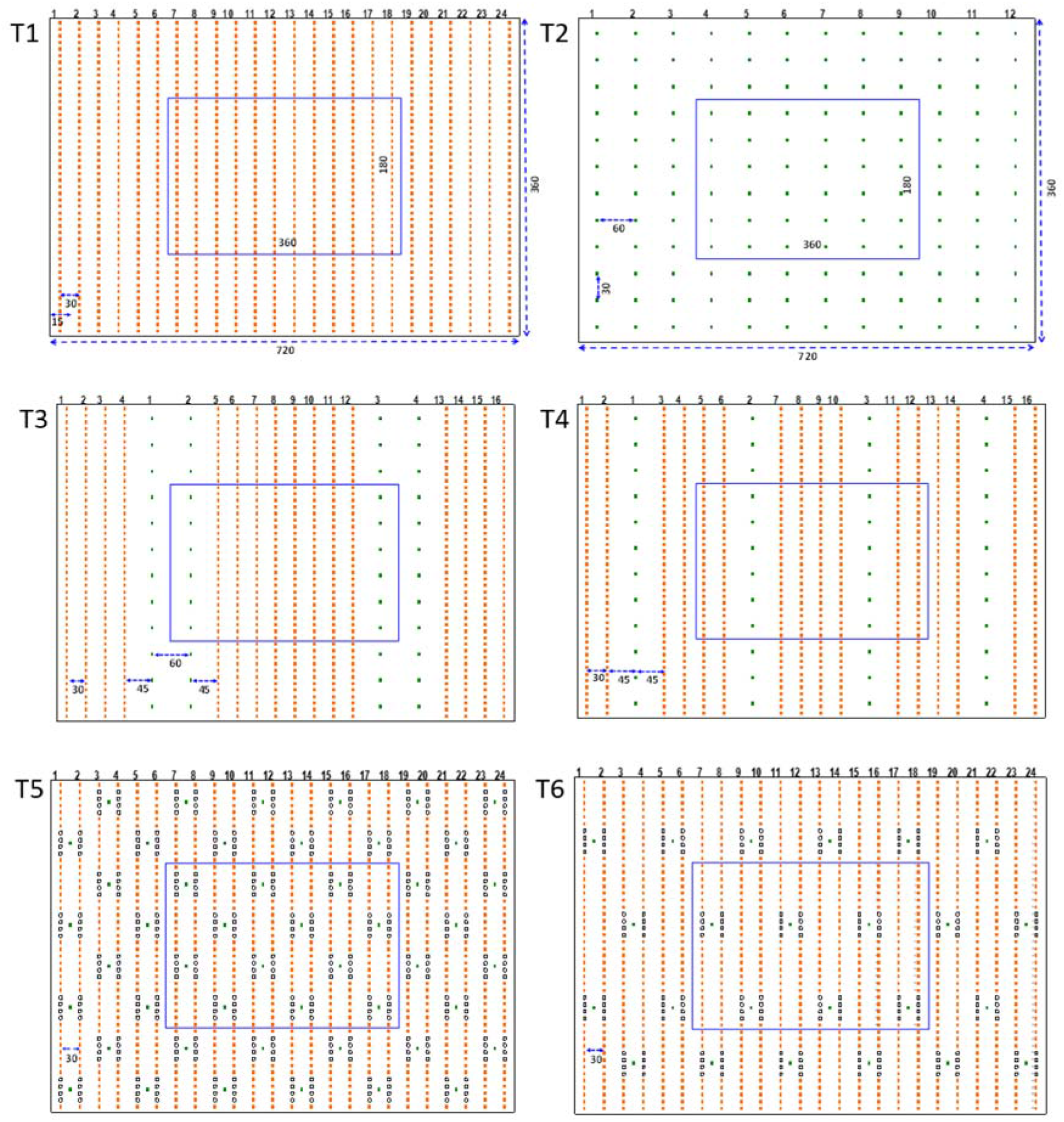
Schematic diagram of field design. Top row: monoculture of FM (T1) and PP (T2). Middle row: 2:8 (PP:FM) row-wise intercropping pattern (T3) and 1:4 (PP:FM) row-wise intercropping pattern (T4). Bottom row: 100% (T5) and 50% (T6) mosaic intercrop design. Number of PP in T6 was reduced to 50% (as compared to T3, T4 and T5; to maintain the planting density similar, FM equivalents were transplanted. In this study, we assumed, 8 FM plants are equivalent to 1 PP plant.

In monocultures, the density of FM was 48 plants per m^2^ and the density of PP was 6 plants per m^2^. We planted 8 times more individuals of FM than PP per area and the total number of plants for FM in monoculture (T1) was 1152 per plot and 288 plants in the net plot area. While, for PP monocrop (T2), the total number of plants was 144 in the total plot and 36 plants in the net plot area. The spacing between FM rows was 30 cm and the distance between FM plants in a row was 7.5 cm. The spacing between PP rows was 60 cm and the distance between PP plants within a row was 30 cm. In intercropping treatments, spacing between PP and FM rows was 45 cm.

Intercropping systems were based on FM monocultures, where eight FM plants were substituted by one PP plant. Row-wise intercropping systems (treatment T3 and T4) were based on previous investigations under rain-fed conditions in Karnataka, India (Ashok *et al*., 2010; Padhi, Panigrahi and Jena, 2010; Mathimaran *et al*., 2020). For T3 (2:8 PP:FM row-wise arrangement), each replicate had thus 48 PP (12 plants x 4 rows) and 768 FM (48 plants x12 rows in each total plot area). T4 (1:4 PP:FM row-wise arrangement) had the identical number of PP and FM plants as T3 but it differed in row arrangement where one row of PP was planted after four rows of FM. Treatment T5 (100% mosaic) consisted of identical numbers of PP and FM plants as T3 and T4, but PP and FM plants were planted within the same row in a mosaic design (Fig. 1b). In treatment T6 (50% mosaic), the number of PP was reduced by 50% and replaced by FM plants. It consisted of 24 PP plants (2 plants x 12 rows) and 960 FM plants. In the 2017-18 field trial at Bengaluru site, FM plants in T5 were not substituted by PP but PP was accidentally added into mosaic design. Therefore, plant density of FM was higher than in the other treatments.

We established the same treatments in the years 2016-17 and 2017-18 except for T6, which was not established in 2017-18 based on results from 2016-17 field trial. While field trials during year 2016-17 had only treatments with biofertilizers, field trials during the year 2017-2018 included treatments with and without biofertilizers (Table 1).

**Table 1.** Intercropping treatments with (AMF + PGPR) and without (none) biofertilizer application were designed and tested at two experimental sites, Bengaluru and Kolli Hills in India. Recommended dose of fertilizer (RDF), and number of FM and PP inside the net plot area are mentioned in the table.

| **Treatment** | **Cropping System** | **PP:FM ratio** | **No. of FM Plant** | **No. of PP Plant** | **Planting system** | **(RDF)** | **Biofertilizer application** |
| --- | --- | --- | --- | --- | --- | --- | --- |
| T1+ | FM | 0:1 | 288 | 0 | Row | 50% | AMF + PGPR |
| T1- | FM | 0:1 | 288 | 0 | Row | 50% | None |
| T2+ | PP | 1:0 | 0 | 36 | Row | 50% | AMF + PGPR |
| T2- | PP | 1:0 | 0 | 36 | Row | 50% | None |
| T3+ | FM+PP | 2:8 | 192 | 12 | Row | 50% | AMF + PGPR |
| T3- | FM+PP | 2:8 | 192 | 12 | Row | 50% | None |
| T4+ | FM+PP | 1:4 | 192 | 12 | Row | 50% | AMF + PGPR |
| T4- | FM+PP | 1:4 | 192 | 12 | Row | 50% | None |
| T5+ | FM+PP | 2:8 (100% PP) | 192 | 12 | Mosaic | 50% | AMF + PGPR |
| T5- | FM+PP | 2:8 (100% PP) | 192 | 12 | Mosaic | 50% | None |
| T6+ | FM+PP | 1:4 (50% PP) | 240 | 6 | Mosaic | 50% | AMF + PGPR |
| T6- | FM+PP | 1:4 (50%PP) | 240 | 6 | Mosaic | 50% | None |

We applied 50% of the recommended dose of fertilizer (RDF) to all plots during sowing of FM seeds, RDF (100%) for PP is 25:50:25 NPK kg ha^-1^ and for FM is 50:40:25 NPK kg ha^-1^. Nitrogen (N) fertilizer was given in the form of Urea (46% N-0P_2_O_5_-0K_2_O, SPIC India Fertilizer Company), Phosphate (P) fertilizer was given in the form of Single Super Phosphate (SSP, 0N-16% P_2_O_5_-0K_2_O, SPIC India Fertilizer Company), and Potash (K) fertilizer was given in the form of Muriate of Potash (MOP, 0N-0P_2_O_5_-60% K_2_O, SPIC India Fertilizer Company).

Biofertilizers consisted of AMF, and plant growth promoting rhizobacteria (PGPR). Two species of AMF inoculants viz. *Rhizophagus fasciculatus* and *Ambispora leptoticha* were selected for FM and PP, respectively, Rhizobium for PP alone, and one PGPR strain (*Pseudomonas* sp. MSSRFD41) for both FM and PP used in this study were as described in Mathimaran et al. (2020). In brief, the two AMF species were multiplied in a vermiculite based carrier material using Rhodes grass (*Chloris gayana*) as a host plant for 40 to 45 days. The harvested dry *A. leptoticha* inoculum, consisting of 24 spores g^-1^ of substrate, was applied at the rate of 5 g per PP seedling (germinated in a polybag, see below) and at ca. 278 kg ha^-1^ (Mathimaran *et al*., 2020). Similarly, *R. fasciculatus*, consisting of 15 spores g^-1^ of substrate was applied at the rate of ca. 444 kg ha^-1^ for FM. The PGPR strains were multiplied in King’s B medium and liquid formulation consisting of 1× 10^9^ CFU per ml of *Pseudomonas* sp. MSSRFD41 (Sekar *et al*., 2018) was applied as seed coating at the rate of 10 ml kg^-1^ seed. Additionally, a band application (along the planting rows) was applied at the rate of 49.5 litres (consisting of 1× 10^9^ CFU per ml) together with farmyard manure (FYM) 7.5 t ha^-1^. The AMFs were obtained from Centre for Natural and Biological Resources and Community Development (CNBRCD), Bengaluru and the PGPR strain was obtained from M. S. Swaminathan Research Foundation (MSSRF), Chennai. In addition, Rhizobium (strain, obtained from Agricultural Station, Amaravati, Andhra Pradesh and liquid formulation was applied as seed inoculum at the rate of 10 ml kg^-1^ PP seeds.

### Pre-germination, sowing of seeds into field, growth period and harvest

Based on an established practice in the area, PP seeds were pre-germinated before planting in polybags (15 x 10 cm) filled with 1.6 kg of a mixture of field soil: FYM:sand (ratio of 15:1:1), and a seed hole of 4 x 1 cm was made at the top(Mathimaran *et al*., 2020). The bottom layer of the seed hole was filled with *A. leptoticha* in vermiculite, two PP seeds coated with rhizobia and PGPR strains were kept above the vermiculite layer and field soil was filled on the top. The seeds were allowed to germinate and grow for 35-45 days. Later, healthy seedlings from these polybags were transplanted into the field during third week of July 2016 for 2016-17 trial, and on first week of August 2017 during 2017-18 field trial. FM seeds were line sown in rows directly into the field immediately after transplanting the PP seedlings, and after germination it was thinned out to maintain the plant density as required in different treatments. FM and PP plants were harvested after 120 and 207 days after sowing, respectively in 2016-17 trial at Kolli Hills, while at Bengaluru site FM and PP were harvested after 127 and 168 days after sowing, respectively. During 2017-18 field trial, FM and PP were harvested at 133 and 245 days after sowing, respectively at Kolli Hills site; at Bengaluru site FM and PP were harvested after 124 and 160 days of sowing.

### Growth and yield parameters

Plant growth parameters such plant height, number of pods, pod weight per plant, number of panicles, grain weight per panicle, straw and grain biomass (both sun dried and oven dried), weight of 1000 FM seeds and 100 seeds of PP were measured after harvesting the plant material in the net plot area. For biomass, plants were harvested row-wise in the net plot area and straw and grains were separated. The sun-dried biomass was determined after drying the straw under the sun for 15 days and 20 days for FM and PP, respectively. Grains were dried under sun for 10 days for PP and FM. A subsample of the sun-dried straw and grain material was oven dried at 80°C for 24 h for calculating the dry matter per row. Biomass per plant was calculated by dividing the row biomass by the number of plants in each row; biomass in tons per ha was obtained by multiplying the row biomass with the number of rows per ha.

### Land equivalent ratio (LER)

The facilitative and competitive interactions between PP and FM in response to the different treatments were calculated using the LER. The LER indicates the efficacy of an intercropping system for using natural resources compared with monoculture (Willey and Osiru, 1972). The baseline for LER is one. If the LER is greater than one, intercropping favours growth and yield of plants, and when it is lower than one, intercropping negatively affects the growth and yield of plants. The LER was calculated as

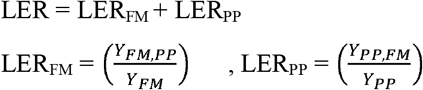

Where Y_FM_ and Y_PP_ are yield of PP and FM in its monoculture, Y_FM,PP_ is yield of finger millet in intercropping, and Y_PP,FM_ is yield of pigeon pea in intercropping.

### Measurement of physiological parameters

Main goal of this study was to test if different spatial arrangements of FM and PP, and the application of biofertilizers affect the water relations and growth of FM. We therefore determined FM leaf water potential at predawn (04:00 to 05:00 hrs) and mid-day (12:30 to 13:30 hrs) towards the end of the field trial during first three weeks on November during 2016-17 and 2017-18. Due to limitation in resources, particularly manpower and time, these measurements were only performed at the Bengaluru site. Both experimental sites received significant amounts of rain till mid of October; therefore, a dry period during November was chosen for measurement (Fig. 2). Leaf water potential (LWP) was measured using a pressure chamber (model 1000, Pressure Chamber Instrument Company, USA). For predawn measurements, leaf samples were collected between 04:00 and 05:00 hours and for midday measurements, leaves were sampled between 12:30 and 13:30 hours. After sampling, leaves were packed into airtight Ziploc bags to avoid water loss; bags were kept in the dark and leaf water potential was measured within 1 – 2 hours after sampling.

**Fig. 2.**
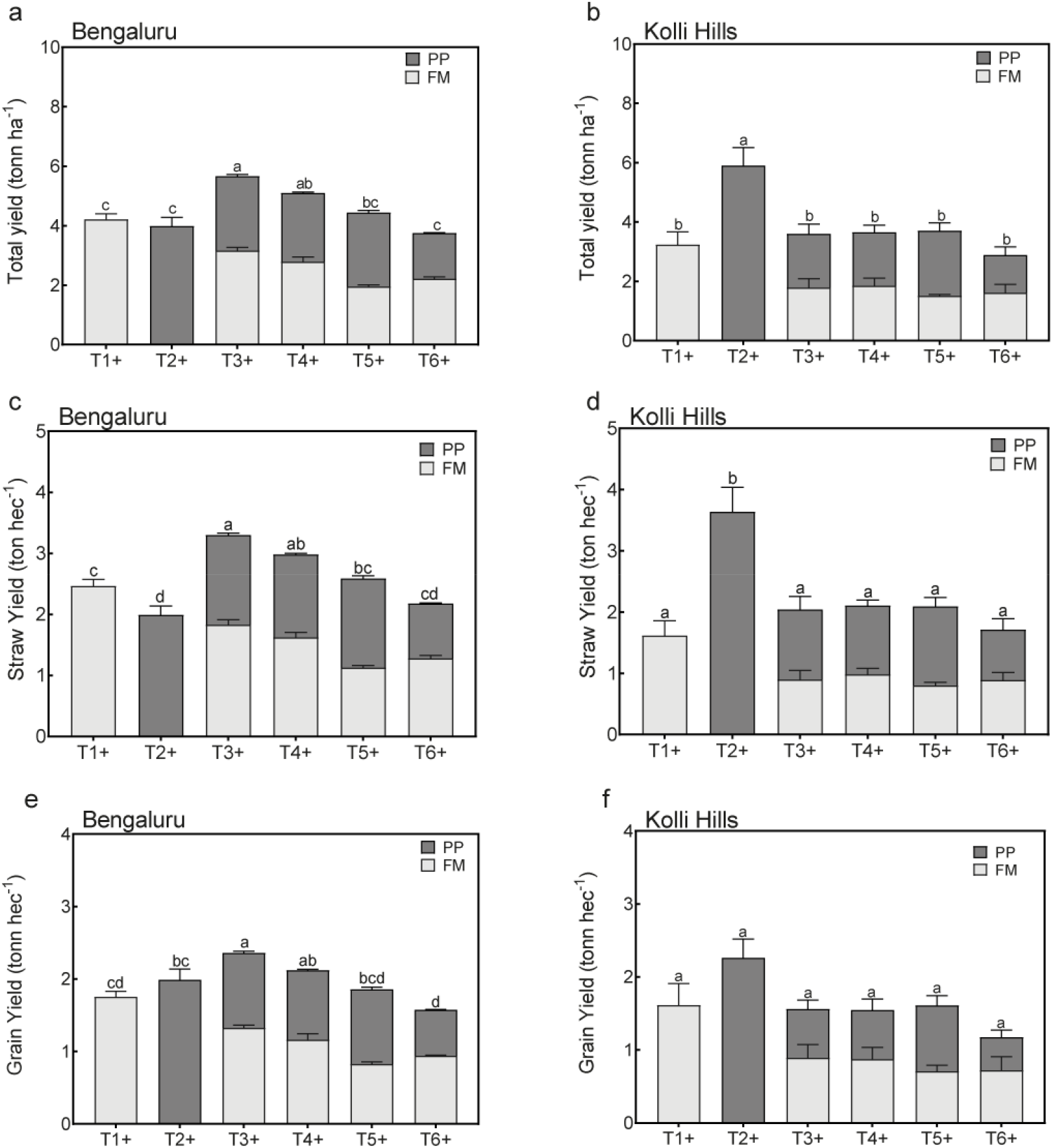
Total biomass, straw and grain biomass of FM and PP at Bengaluru (Fig. 2a, 2c & 2e) and Kolli Hills (Fig. 2b, 2d & 2f) during year 2016/17. Bars represent the average of four replicates with standard error of mean. One-Way ANOVA followed by Tukey’s test (posthoc test) was used for the combined biomass of FM and PP, separately for each site, and values with same letters are not significantly different from each other at p>0.05.

### Statistical analysis

Analysis of yield data and LWP from field trials was carried out using GraphPad Prism software (version 7.0 for Mac OS X, GraphPad Software, La Jolla California USA). Data are expressed as mean ± standard error of mean (SEM). Tukey’s test was used for post hoc multiple treatment comparison following one-way ANOVA or multifactor ANOVA using general linear models. The criterion for significance was p<0.05.

## Results

### Total biomass, straw and grain yield per hectare and LER

Intercropping and the spatial arrangement of the intercropping partners had a significant effect on the total biomass yield per hectare at the Bengaluru site in 2016-17 (Fig. 2, Table S1). In particular, the treatment T3+ produced significantly more biomass per hectare than monocultures of the constitutive crops or other spatial arrangements at Bengaluru in 2016-17. Likewise, treatment T3+ resulted in higher yields for straw and grain as compared to the other treatments in 2016-17 at Bengaluru site (Fig. 2, Table S1). For the intercropping treatments, total biomass yield, straw yield and grain yield all declined from the T3+ to T6+. The results differed at the Kolli Hills site, where in 2016-17 PP (T2+) produced the highest yields for total biomass, straw and grain and where FM (T1+) and the different intercropping treatments produced slightly lower yields with no significant differences among each other (Table S1). In summary, in 2016-17 we found a strong positive intercropping effect for total biomass yield, straw yield and grain yield at Bengaluru site, where the intercropping effect were strongest in the 8:2 row-wise spacing. In contrast, no yield improvements by intercropping irrespective of the spatial arrangement were observed at the Kolli Hills site.

These observations are also reflected in LER values at Bengaluru site, where values for total biomass were greater than one for T3+, T4+ and T5+ and where T3+ had the highest LER value. Similarly, for straw biomass, T3+ had higher LER values than T4+, T5+ and T6+. For grain biomass LER values were greater than one for the T3+ and T4+ treatment, equal to one for T5+ and less than one for T6+ (Fig. 3). At Kolli Hills LER values for all treatments were less than one (Fig. 3).

**Fig. 3.**
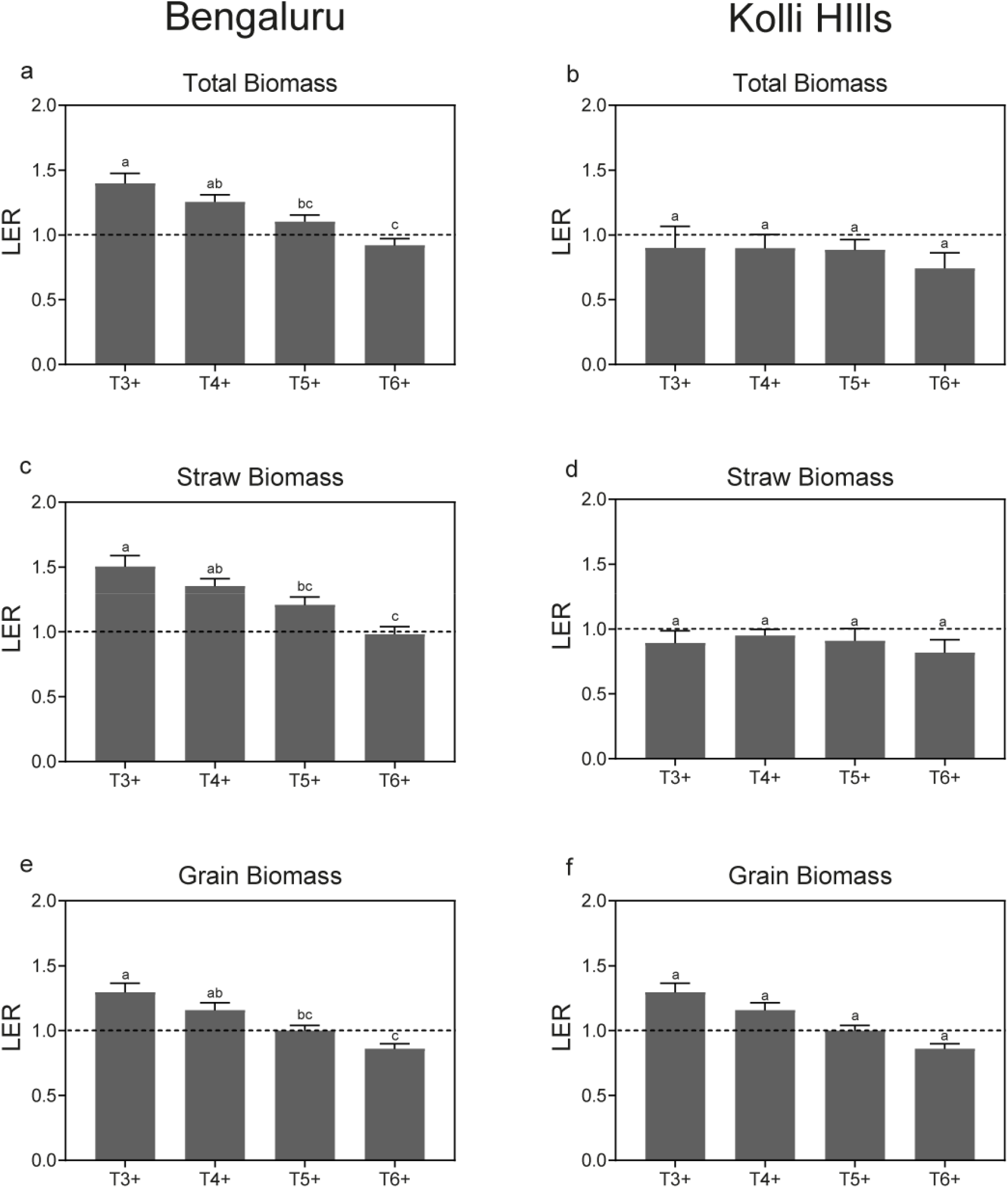
Land equivalent ratio (LER) in different intercropping treatments during 2016-17 at Bengaluru (Fig. 3a, 3c & 3e) and Kolli Hills (Fig. 3b, 3d & 3f) site. Bars represent the average of four replicates with standard error of mean. Tukey’s test (one-way ANOVA) was used for multiple comparison, separately for each site, and values with same letters are not significantly from each other different at p>0.05.

In 2017-18, intercropping and the spatial arrangement of the intercropping partners also had a strong and significant effect on the total biomass yield, straw yield and grain yield at Bengaluru site (Fig. 4). As in 2016-17 the treatment T3-and T3+ produced significantly more biomass per hectare than monocultures of the constitutive crops or other spatial arrangements when compared to the respective treatments with and without biofertilizer. Importantly, the application of biofertilizers enhanced the total biomass yield, straw yield and grain yield in all treatments and this effect was consistent irrespective of experiment site, mono or intercropping (Table S2). At Kolli Hills, we also found significant treatment effects (Fig. 4). However, intercropping treatments did not produce higher yields for total biomass and straw than any of the other treatments with or without biofertilizer. Yet, treatment T5+ was equal in total biomass yield than the most productive monoculture (T2+). For grain yield FM monoculture exceeded the productivity of PP (Fig. 4f) and in intercropping T3-, T3+ and T5+ grain yield was similar to monoculture of FM with or without biofertilizer. The effects of biofertilizers on total biomass yield, straw yield and grain yield that we detected at the Bengaluru site were also observed at the Kolli Hills site and this effect was again consistent across all treatments (Fig. 4, Table S2). We did not find a significant interaction between treatment and biofertilizers nor a significant three-way interaction between treatment, biofertilizers, and site. However, as indicated above, the effects of biofertilizers at Kolli Hills resulted in total biomass yield, straw yield and grain yield that were of the same magnitude in some intercropping treatments as the highest yield in the corresponding monocultures (e.g. T5+ for total biomass yield, and straw yield, and T3+ and T5+ for grain yield) (Fig. 4). In summary, in 2017-18 we found a strong positive intercropping effect for total biomass yield, straw yield and grain yield at Bengaluru site. In Kolli Hills, no such intercropping effect was found. Importantly, biofertilizers improved the yields of crops in both sites and independently of treatment. Despite the nonsignificant biofertilization – treatment interaction, intercropping treatments at Kolli Hills showed yet a trend to be more enhanced through biofertilizers than monocultures to an extent that they produced similar yields than the most productive monoculture, which we did not observe without biofertilizers.

**Fig. 4.**
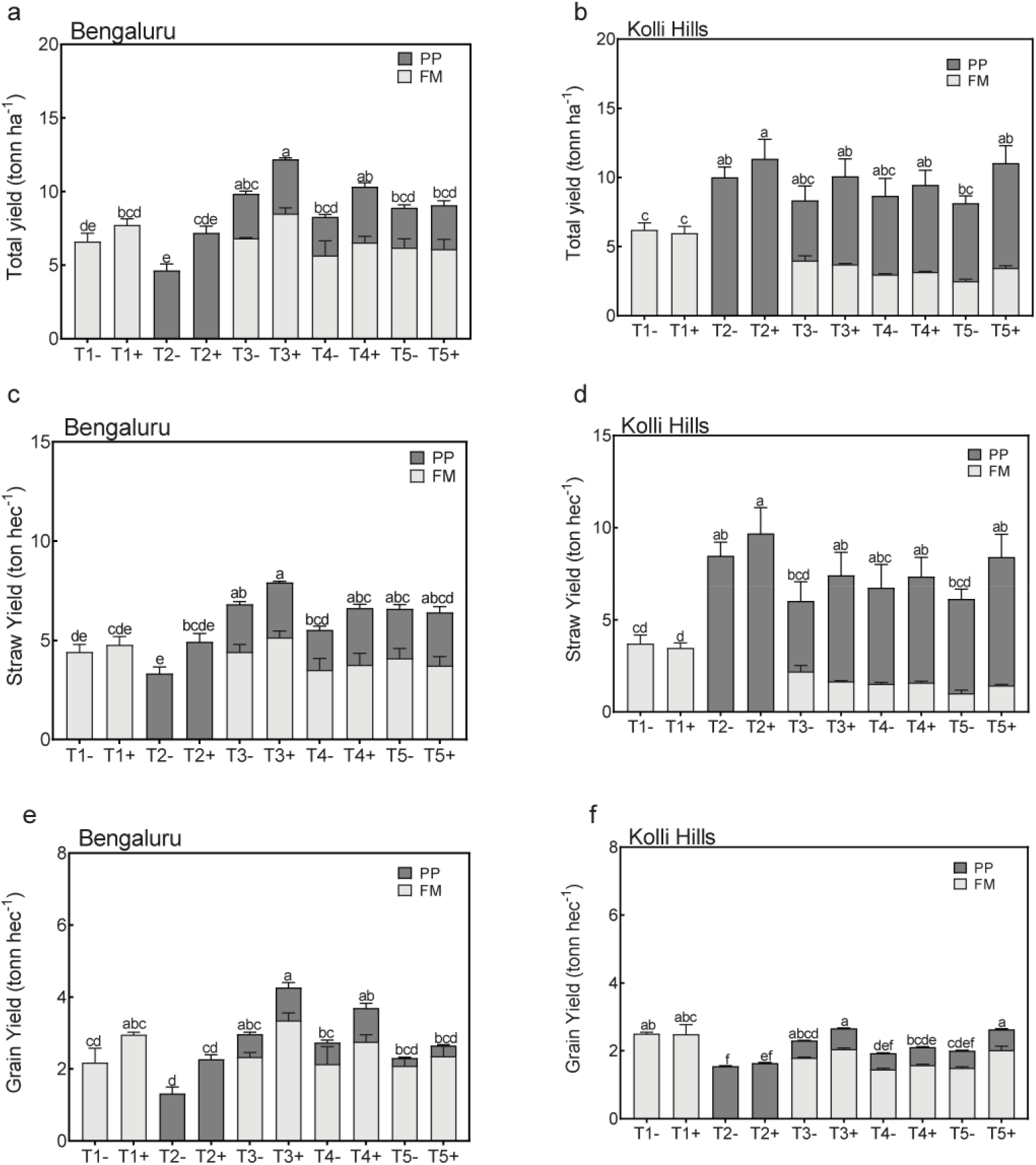
Total biomass, straw and grain biomass at Bengaluru (Fig. 4a, 4c &4e) and Kolli Hills (Fig. 4b, 4d & 4f) during year 2017-18. Bars represent the average of four replicates with standard error of mean. One-way ANOVA followed by Tukey’s test (posthoc test) was used for the combined biomass of FM and PP, separately for each site, and values with same letters are not significantly different from each other at p>0.05.

These observations were confirmed by LER values for 2017-18 at both sites (Fig. 5). LER was greater than one at the Bengaluru site for all treatments. Also, LER values at the Bengaluru site were largest for T3+ and declined in the other treatments. Biofertilizers had a negative effect on LER values in all spatial arrangements at the Bengaluru site. At Kolli Hills, LER values in treatments without biofertilizers were either equal to or less than one. Biofertilizers increased, however, the LER values in all spatial arrangements to values of one or greater than one and the largest values were observed for T3+ and T5+.

**Fig. 5.**
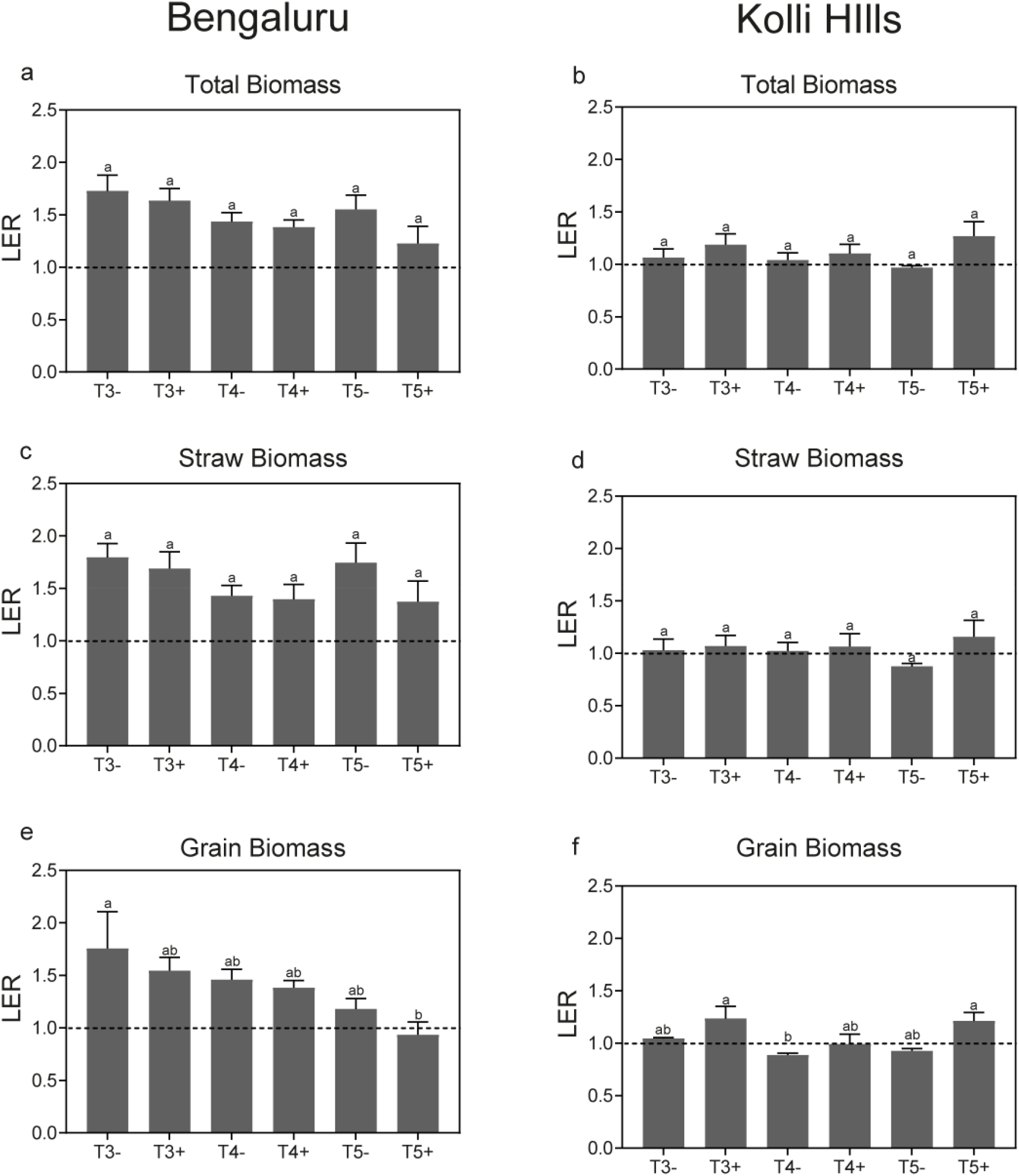
Land equivalent ratio (LER) of total grain yield in different intercropping treatments during 2017-18 at Bengaluru (Fig. 5a, 5c & 5e) and Kolli Hills (Fig. 5b, 5d & 5f) site. Bars represent the average of four replicates with standard error of mean. Tukey’s test (one-way ANOVA) was used for multiple comparison, separately for each site, and values with same letters are not significantly different from each other at p>0.05.

### Per plant biomass yield of PP and FM

We found a significant effect of the intercropping treatments on total biomass per plant, total straw yield per plant and total grain yield per plant of PP and FM at the Bengaluru site but not in Kolli Hills in 2016-17 (Fig. 6, Table S3 & S4). At Bengaluru, total biomass per plant in FM was highest in the monoculture (T1+), the 2:8 treatment (T3+) and the 1:4 treatment (T4+). The biomass of the individual plants was significantly reduced in the mosaic treatments (T5+ and T6+) compared to monoculture (T1+) and row-wise intercropping (T3+ and T4+, Fig. 6a). PP showed highest total biomass in the mosaic treatment T6+, followed by other intercropping treatments and lowest biomass in the monoculture T2+ (Fig. 6c). At Kolli Hills, total biomass per plant in PP and FM did not differ significantly among treatments (Fig. 6b &6d). However, the trend was similar to the Bengaluru site where FM showed a reduction in biomass in mosaic treatments while PP showed an increase in biomass in mosaic treatments.

**Fig. 6.**
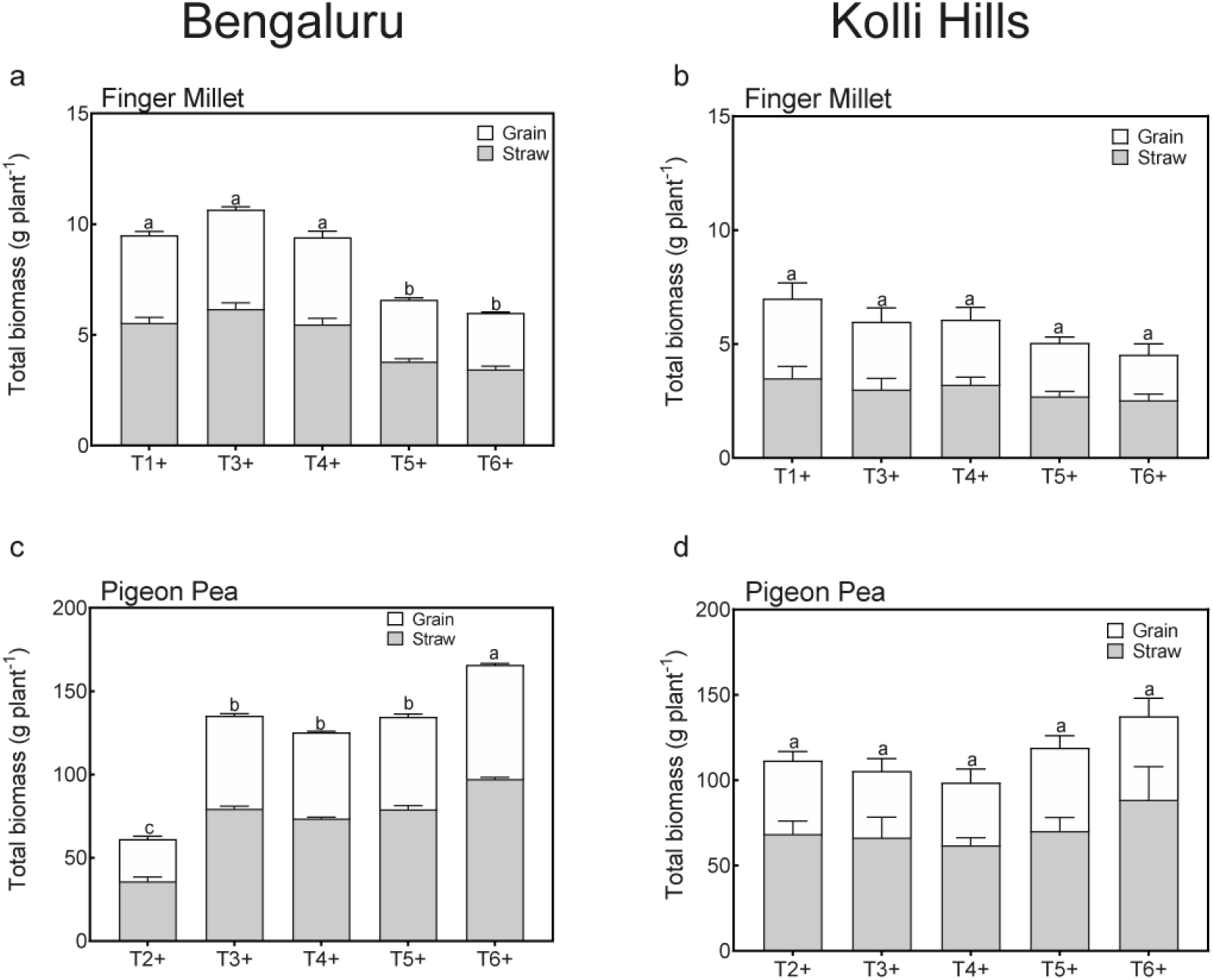
Total biomass per plant of FM and PP at Bengaluru site (Fig. 6a & 6c) and Kolli Hills site (Fig. 6b& 6d) during 2016-17 field trial. Bars represent the average of four replicates with standard error of mean. One-way ANOVA followed by Tukey’s test (post hoc test) was used for the combined biomass of grain and straw, separately for each site, and values with same letters are not significantly different from each other at p>0.05.

In 2017/18 we also found a significant treatment effect on the total biomass, straw yield and grain yield of FM and PP at the Bengaluru site but only for PP at Kolli Hills (Fig. 7, Tables S5 & S6). At the Bengaluru site, total biomass of FM plants in T3+ was significantly larger than total biomass of plants in treatments T1-, T1+ and T4-. Total biomass of PP plants were largest in T3+ and T5+ compared to T2-, T2+, and T4-. At Kolli Hills total biomass per plant in FM did not show any significant difference among intercropping and monoculture. For PP, in contrast, total biomass per plant was largest in treatments T4+ and T5+ compared to T2-and T2+ (Fig. 7d).

**Fig. 7.**
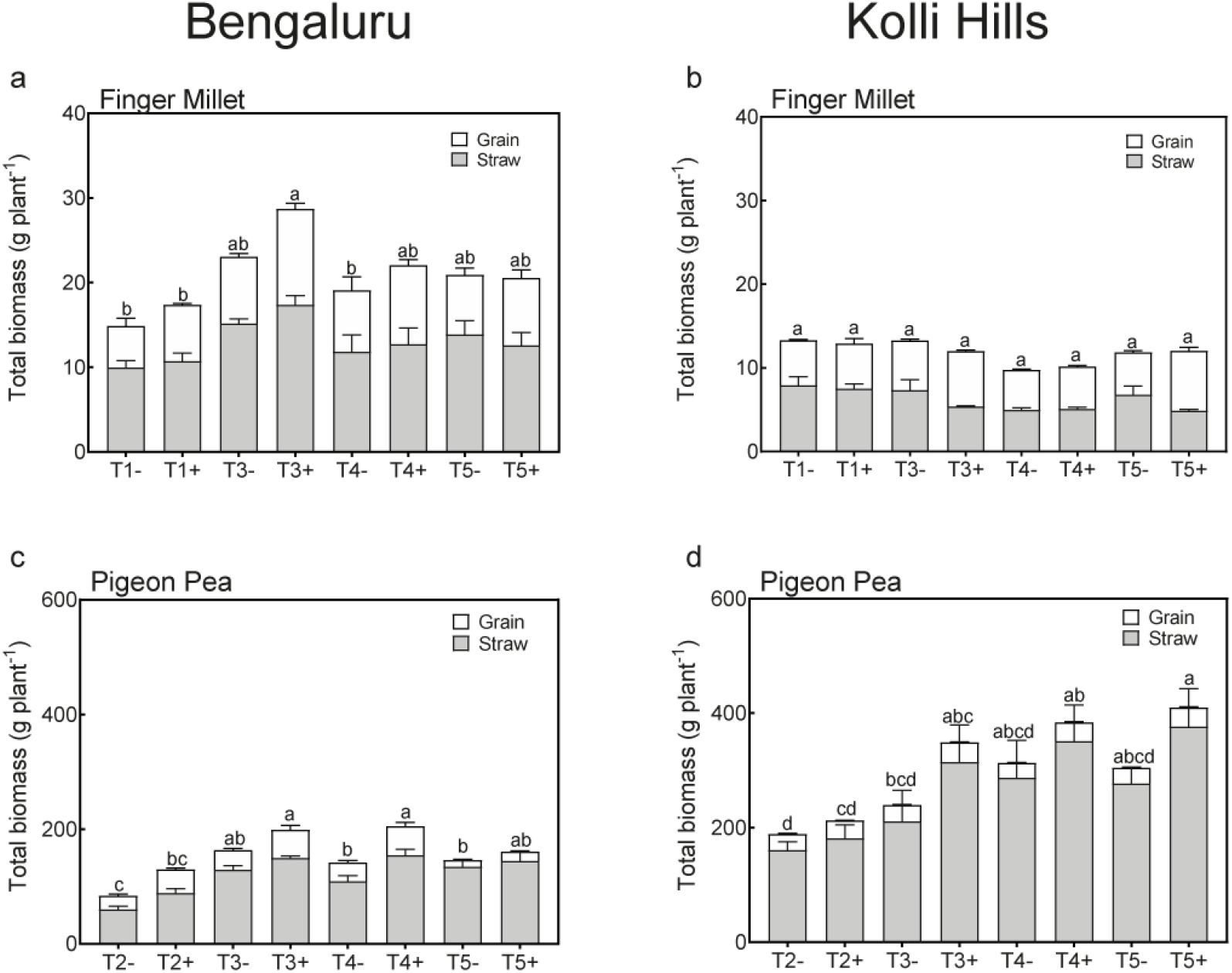
Total biomass per plant of FM and PP at Bengaluru site (Fig. 7a & 7c) and Kolli Hills site (Fig. 7b & 7d) during 2017-18 field trial. Bars represent the average of four replicates with standard error of mean. One-way ANOVA followed by Tukey’s test (post hoc test) was used for the combined biomass of grain and straw, separately for each site, and values with same letters are not significantly different from each other at p>0.05.

A two-way ANOVA analysis was performed to test the effects of spatial arrangement and biofertilization on per plant yield (Table S7 & S8). At both sites in 2017-18 FM yield did not show any significant effect of biofertilizer application. However, PP showed a strong significant effect of biofertilization at the Bengaluru site, and at the Kolli Hills site the effect was marginally significant. At both sites, the effect of biofertilization did not differ among treatments due to spatial arrangement of the component plants in an intercropping system.

### Water relations of PP and FM in intercropping treatments

Measurements of the predawn leaf water potential (LWP) were done FM leaves at Bengaluru site to evaluate the effect of spatial arrangement and biofertilizer application on the water relations of FM in different intercropping treatments (Fig. 8, Table S9). In 2016-17 in week 1 of the measurements (1^st^ week of November 2016), FM in treatment T1, which is the monoculture treatment, had the most positive values (−0.70 MPa). FM in the mosaic treatment T5+ had the lowest predawn LWP of -2.5 MPa, which is significantly lower than in the row-wise intercropping treatment (T3+, -0.95 MPa). In week 2 (2^nd^ week of November 2016), FM in monoculture (T1+) maintained a significantly higher predawn LWP of -1.15 MPa than in any other intercropping treatment (Fig. 8a). At week 3, (3^rd^ week of November 2016) FM in treatments T4+ and T5+ were dead (desiccated & drooped), while FM in T3+ and T6+ showed a significantly lower LWP of -1.89 and -1.90 MPa than FM in monoculture (−1.34 MPa).

**Fig. 8.**
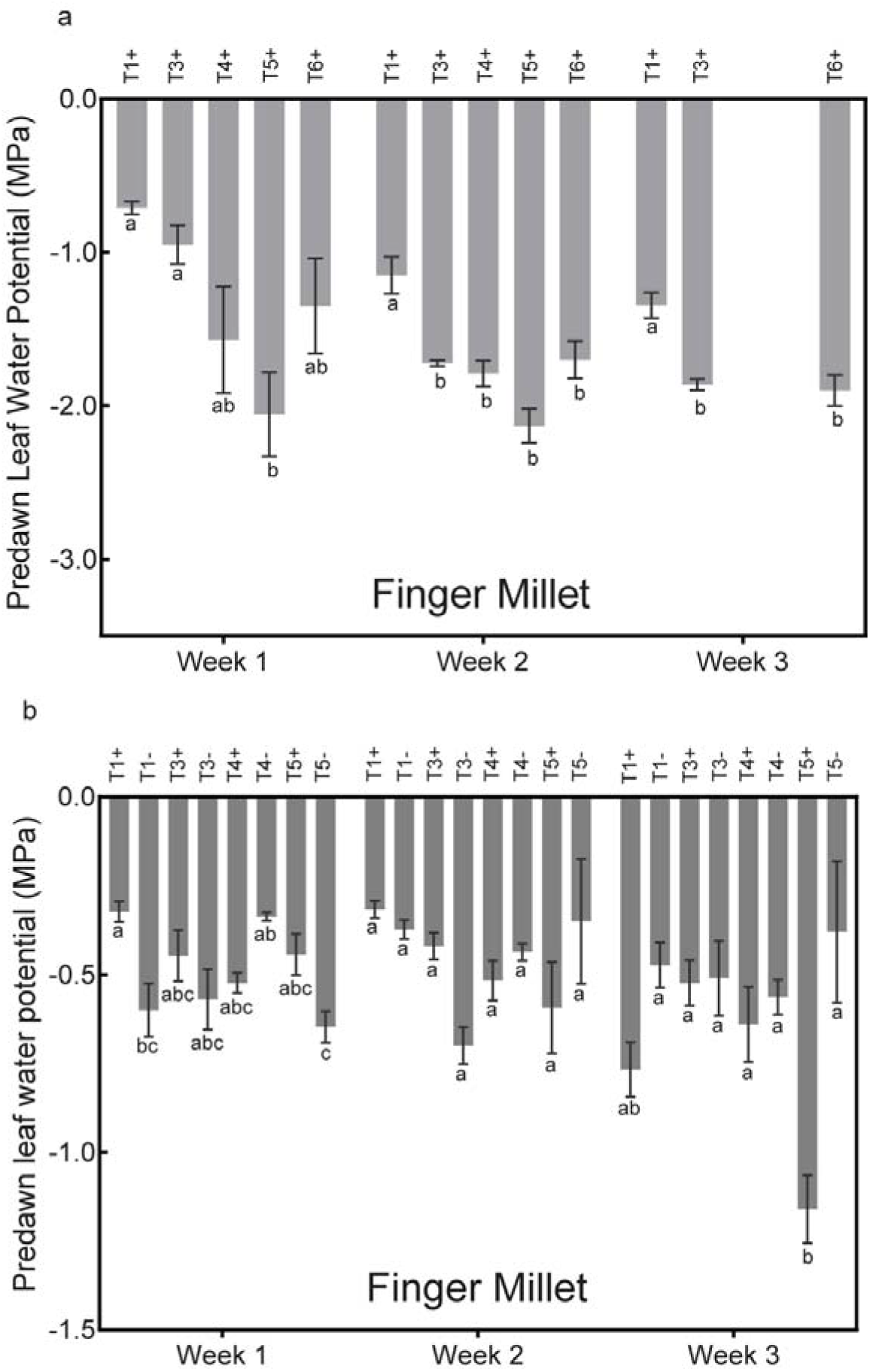
Predawn leaf water potential of FM in different intercropping treatments during 2016/17 (Fig. 8a) and 2017-18 (Fig. 8b) field trial. Weeks represent first, second and third week of November in 2016 and 2017, during which measurement was done. Bars represent the average of four replicates with standard error of mean. Tukey’s test (One-way ANOVA) was used for multiple comparison and values with same letters are not significantly different from each other at p>0.05.

In 2017-18 at week 1 (1^st^ week of November 2017), predawn LWP of FM in monoculture with biofertilizer (T1+) had values of -0.32 MPa which is significantly more positive than FM in monoculture without biofertilizer (T1-) (−0.60 MPa) or any of the intercropping treatments (Fig. 8b). Later, FM did not show any significant difference in LWP compared to the other intercropping treatments. Interestingly, treatments without biofertilizer showed lower values for LWP as compared to the respective treatments with biofertilizer. The biofertilizer application did not have a significant effect on LWP of FM, but intercropping treatments showed a strong significant effect (Table S9). Effect of biofertilizer showed significant interaction with intercropping treatments, as we observed in fig. 8, treatments T1-, T1+, T5-and T5+ consistently showed a large difference in LWP of FM with or without biofertilizer.

## Discussion

The results obtained from the field trials during 2016-17 and 2017-18 showed that intercropping can improve the straw and grain yield in PP–FM intercropping compared to the respective monocultures but that intercropping effects vary depending on the site characteristic such as climate and soil type as well as crop variety. Spatial arrangement of component plants affected the total, straw and grain biomass in intercropping treatments, but this effect also varied across sites. The results from 2017-18 clearly demonstrated a positive effect of biofertilizer on biomass yield, and this effect was irrespective of site, spatial arrangement, mixed or monoculture. Despite the positive effect of intercropping and biofertilization on FM and PP yield, water-relations of FM were not enhanced in the intercropping treatments or by biofertilizers. Most likely this is due to interspecific competition for soil moisture in top soil layer between PP and FM. On the basis of these results, we propose that intercropping and the application of biofertilizer both enhance the yield of cropping systems and effects on yield if intercropping and biofertilization are applied in combination. However, the spatial arrangement of component crops is a key factor that affects the productivity of the involved intercropping partners.

### Is PP – FM intercropping beneficial over monocropping?

The yield advantage in intercropping systems is typically assigned to resource sharing and facilitation (Loreau and Hector, 2001; Li *et al*., 2014; Duchene, Vian and Celette, 2017). Resource complementarity reduces the niche overlap and competition between two species and allow crops to uptake greater range of resources than the sole crops. Ghanbari et al. (2010) reported resource complementarity in maize-cowpea intercropping systems, where intercropping increased the light interception, reduced evaporation, and improved soil moisture conservation compared to maize sole crops. In most cases, facilitation occurs through increased availability of soil resources such as water and nutrients (Jensen, Carlsson and Hauggaard-Nielsen, 2020). Intercropping systems with legume species (such as PP in this study) can increase agricultural productivity through providing increased nitrogen availability through N_2_ fixation, and are therefore used very frequently in intercropping systems (Hauggaard-Nielsen and Jensen, 2005; Altieri, Funes-Monzote and Petersen, 2012). Nonlegumes (such as cereals) in an intercropping system with legume plants may obtain additional N released by legumes into soil, and legumes can contribute up to 15% of the N in intercropped cereals (Li et al., 2007; Zuo et al., 2004).

According to our expectation, we found at the Bangalore research site, that intercropping treatments (T3+ and T4+) produced higher yields (Fig. 2 & 4) than monocultures in both growing seasons. In contrast, in Kolli Hills, there was no significant effect of intercropping in the 2016-17 season. In 2017-18 we also did not observe strong intercropping effects but yields in some intercropping treatments (T5+) were as high as the highest yields in the monocrop. Accordingly, LER values were above one in Bangalore in both years but below one in Kolli Hills in 2016-17 and near zero or above in 2017-18. This illustrates that intercropping effects depend on the climate of the growing season and soil type at the experiment site. The total rainfall in Kolli Hills in 2017 was 1690 mm, while the 2016 growing season was shaped by a severe drought with a total rainfall of only 281.7 mm. Additionally, both locations differ in their soil properties. At the Bangalore site, the soil is an Alfisol with 67.8% sand, 7.7% silt, 25.2% clay, C_org_ 0.5% and a pH of 4.8. At the Kolli Hills site, the soil type is a Vertisol with 33.2% sand, 30.0% silt, 36.8% clay, C_org_ 0.8% and a pH of 5.2 (see Mathimaran et al., 2020, supplimentary data). The relationship between crop yield and soil depends on complex interactions between physio-chemical properties of soil and other climatic factors (Stenberg, 1998). Juhos et al. (2015), using a multivariate statistical approach, show that in droughty years the sodification, salinization, soil texture and nutrient content determined the yield, while in humid years soil organic matter and nutrient content were the main determining factors for crop yields. Our results indicate that the low amount of rainfall and inherent soil properties could be the factor which caused different intercropping effect at the two sites and between the two growing seasons (Fig. 2).

### Effect of spatial arrangement on yield in PP -FM intercropping

At the Bengaluru site, straw and grain yield (per hectare) showed that row-wise intercropping treatments produced higher yield than mosaic during 2016-17 and 2017-18 field trial. The results from Kolli Hills were inconsistent, perhaps because of rainfall and soil properties. Effects of the spatial arrangements can be explained by intra-and interspecific competition, as illustrated when data are expressed per plant biomass (Fig. 6 & 7). Results from Bengaluru clearly indicate that PP benefits in terms of per plant biomass in intercropping treatments likely due to reduction in intra-specific competition that PP faces in monoculture. In contrast, FM faces higher inter-specific competition in mosaic treatments, which leads to a reduction in per plant biomass in mosaic treatments (T5+ and T6+). The field trial results from Kolli Hills, however, do not show any significant effect of spatial arrangement of plants on per plant biomass in PP and FM during 2016-17 trial (Fig. 8b & 8d). During 2017-18, only PP showed a significant increase in per plant biomass in intercropping treatments T4+ and T5+ compared to monoculture treatments T2-and T2+. The effect of spatial arrangement on FM per plant biomass was not significant and it was consistent during both years at Kolli Hills.

The results of this study show consistently that PP growth is favoured in intercropping systems due to reduction in intra-specific competition, while FM faces higher inter-specific competition in mosaic intercropping than in row-wise intercropping. This effect is modulated by the variety (different varieties of PP were grown at Bangalore and Kolli Hills research site) of intercropped PP, soil quality and local weather. There are several factors, such as light, soil moisture and nutrient, that affect the yield of each component crop in intercropping (Bedoussac *et al*., 2015). The difference in penetration of light into canopy is considered to be a key factor affecting photosynthesis and ultimately growth and yield (Gwathmey and Clement, 2010; Kaggwa-Asiimwe, Andrade-Sanchez and Wang, 2013). In our study, the reduction in light availability to relatively short FM plants standing next to taller PP plants in the mosaic intercropping treatments T5 & T6 (see supplementary data) could be a factor impacting growth, since in all row-wise intercropping designs PP and FM rows are well spaced to avoid a shading effect, which is not the case in the mosaic design. Similar results were reported by Martin and Snaydon (1982) and Dubey et al. (1995), who reported highest yield for barley/beans and sorghum/soybean in row-wise intercropping than mosaic (mixed within rows), respectively.

The intercropping designs tested in this study illustrate that the row-wise intercropping treatment T3+ (2:8 with biofertilizer) performed consistently better than the other arrangements, which is due to the release of intra-specific competition. Effects of the spatial arrangement of component plants in intercropping have been shown to be species specific. Chen et al. (2004), Lauk and Lauk (2008) and Aynehband et al. (2010) have shown mixing of component plant within rows (mosaic pattern) to be the best arrangement for barley/peas, maize/soybean and maize/amaranth, respectively. In contrast, Martin and Snaydon (1982) and Dubey et al. (1995) reported higher yields for barley/beans and sorghum/soybean sown in alternate rows than mixed within rows, respectively. Interspecific competition could occur when two species are planted together, and such competition could lead to decrease in plant growth and yield (Jensen, 1996). In a cereal-legume intercropping system there is a significant number of days for overlapping growth period, and interspecific competition between component crops could lead to a decrease in yield (Clément *et al*., 1992; Oljaca *et al*., 2000; Karasawa and Takebe, 2012); therefore, spatial arrangement between the plants needs to be carefully optimized. In this study, PP had a head start of 45 days (polybag transplantation) compared to FM, which provided PP a competitive advantage to acquire more resources (light, nutrients and water) through its well-established root network, and FM may face, additionally, shading effect due to tall PP plants.

### Effects of biofertilizers

In the 2017-18 field trial, at both experimental sites, the effect of biofertilizer application was positive and showed an increase in total yield (Fig. 7). The positive effect of biofertilization did not differ among intercropping treatments with different spatial arrangements (Table S7 & S8). The effect of biofertilization was, however, specific to each component plants in the PP–FM intercropping system. Total biomass and straw yield per plant in FM was not significantly affected by biofertilization, but grain yield was significantly increased (Table S5) similar to observations made in Mathimaran et al. (2020). In the case of PP, the effect of biofertilization was significant on total biomass, straw and grain yield per plant. The results of this study are in agreement with findings of Mäder et al. (2011) who reported that combined application of AMF and PGPR improves grain yield. Previous studies (Reddy, 2012; Mathimaran *et al*., 2017) have shown that better phosphorus uptake and crop tolerance to biotic and abiotic stresses via PGPR are among the most common mechanisms through which biofertilizers improve crop growth. The increase in grain yield in both component plants (FM and PP) in intercropping was the result of an increased number of panicle and grain weight per panicle in FM and number of pod and pod weight per plant in PP (see supplementary data). Since the process of pod and panicle formation is influenced by light availability, nutrients and soil moisture (Härdter and Horst, 1991), the yield improvement in row-wise intercropping could be attributed to efficient utilization of nutrients through the applied biofertilizers.

### Effect of intercropping and biofertilizers on water relations of FM

In this study, the water relations (predawn LWP) of FM decreased significantly in mosaic treatments as compared to row-wise and monoculture treatments (Fig. 8a & 8b). The trend in predawn LWP (Fig. 8a & 8b) can also be compared with the trend in biomass production per plant (Fig. 6a & 7a), therefore, competition for water could be the limiting factor here which influenced the yield and effectiveness of intercropping treatments at Bengaluru site. Our results suggest that there exists an important degree of below-ground competition for water between PP and FM, and the facilitative effect of bioirrigation is suppressed. Similar results have been reported by Ludwig et al. (2004). They found that HL performing trees extracted a significant amount of water from the topsoil layer that resulted in lower LWP in understorey grasses; however, grasses were able to absorb soil moisture released by tree due to HL.

One of our objectives was to find out if CMN can facilitate the transfer of bioirrigated water from PP to FM and improve the water-relations of FM in intercropping treatments. The results from the 2017-18 field trial showed that CMN did not affect the water relations (predawn LWP) of FM in intercropping treatments. However, at week 1 and 2 (first and second week of November 2017) FM in T3+ had higher, but not significant, LWP than T3-. Similarly, FM in monoculture treatment showed a higher (less negative LWP) with CMN than without CMN (Fig. 8b). Since, we observed similar effects of CMN in both monoculture and 2:8 row-wise intercropping, we cannot assign this to bioirrigation. The effect of CMN changed over time, and at week 3 (third week of November 2017) treatments T1+, T3+, and T5+ (with CMN) had a lower LWP than T1-, T3-, and T5-(without CMN). The effect of different treatments, biofertilization and times (weekly measurement) had significant interaction with each other (Table S9).

In this study, we could not find out if the positive intercropping effect by CMN was due to bioirrigation. The average hyphal spread rate of *Glomus* species is 0.7 – 0.8 mm per day (Jakobsen, Abbott and Robson, 1992), We did not check for the spread of CMN between PP and FM, but it is possible that the AMF introduced with the biofertilizer could not cover the distance of 45 cm between PP and FM in intercropping treatments and thus, a potential facilitative effect of bioirrigation through CMN was not observed.

## Conclusions

This study, to our best knowledge, for the first time shows that spatial arrangement of component crops affects the competition for topsoil moisture between crops in rainfed areas Furthermore, we clearly demonstrate that intercropping has a positive effect on total yield of PP and FM but this effect varies across the sites based on site characteristics such as soil type and weather. In conclusion, the answers to our three research questions are as follows: (i) the spatial arrangement of intercropping partners does affect the straw and grain yield in a FM – PP intercropping system, and the optimal spatial arrangement for PP – FM intercropping system depends on geographic location (local weather conditions) and plant variety. In general, the row-wise treatment (T3+) resulted in better yields than the mosaic treatments at Bengaluru site, while at Kolli Hills site in 2017-18, both row-wise treatment (T3+) and mosaic treatment (T5+) performed equally well. Most importantly, (ii) we show that the application of biofertilizer promotes yield in intercropping system, and the spatial arrangement of component plants do not affect the effect of biofertilization. The effect of biofertilization is mainly due to the promotion of PP. We further show that (iii) the spatial arrangement of plants is a key factor that affects the competition for topsoil moisture between PP and FM.

Further research with different varieties of PP, and different spatial arrangement including the planting distance between PP and FM will provide crucial information to design bioirrigation based intercropping models for rainfed areas in semiarid tropics.

## Ethics Approval

Not relevant for this work.

## Consent to participate

Not relevant for this work.

## Consent for publications

Not relevant for this work

## Data availability

All data generated or analyzed during this study are included in this published article.

## Code availability

Not relevant for this work.

## Authors’ contribution

**Conceptualization:** AK, NM, PVR, IOK, TMN, PM, TB, DS, JS

**Methodology:** AK, NM, PVR, TMN, PM, TB, DS, JS

**Formal Analysis:** DS

**Resources:** AK, NM, PVR, RR, IOK, TMN, DJB,

**Data curation:** DS, JS, YP, KR, NM, IOK, MBN, BNC, SMS, TMN

**Writing – original draft preparation:** DS

**Writing – review and editing:** DS, NM, JS, PVR, YP, KR, RR; IOK, TMN, MBN, BNC, SMS, DJB, PM, TB, AK

## References

Altieri, M. A., Funes-Monzote, F. R. and Petersen, P. (2012) ‘Agroecologically efficient agricultural systems for smallholder farmers: Contributions to food sovereignty’, Agronomy for Sustainable Development, 32(1), pp. 1–13. doi: 10.1007/s13593-011-0065-6.

Ashok, E. G. et al. (2010) ‘Augumenting production and profitability of finger millet+pigeonpea intercropping system’, Environment & Ecology, 28(1), pp. 28–33.

Augé, R. M. et al. (2001) ‘Moisture retention properties of a mycorrhizal soil’, Plant and Soil, 230(1), pp. 87–97. doi: 10.1023/A:1004891210871.

Aynehband, A., Behrooz, M. and Afshar, A. H. (2010) ‘Study of intercropping agroecosystem productivity influenced by different crops and planting ratios’, American Eurasian Journal of Agricultural and Environmental Science, 7(2), pp. 163–169.

Bedoussac, L. et al. (2015) ‘Ecological principles underlying the increase of productivity achieved by cereal-grain legume intercrops in organic farming. A review’, Agronomy for Sustainable Development, 35(3), pp. 911–935. doi: 10.1007/s13593-014-0277-7.

Bogie, N. A. et al. (2018) ‘Intercropping with two native woody shrubs improves water status and development of interplanted groundnut and pearl millet in the Sahel’, Plant and Soil. 435, pp. 143– 159. doi: 10.1007/s11104-018-3882-4.

Brooker, R. W. et al. (2015) ‘Improving intercropping: a synthesis of research in agronomy, plant physiology and ecology’, New Phytologist, 206(1), pp. 107–117. doi: 10.1111/nph.13132.

Brooks, J. R. et al. (2006) ‘Hydraulic redistribution in a Douglas-fir forest: Lessons from system manipulations’, Plant, Cell and Environment, pp. 138–150. doi: 10.1111/j.1365-3040.2005.01409.x.

Burgess, S. S. O. (2011) ‘Can hydraulic redistribution put bread on our table?’, Plant and Soil, 341(1– 2), pp. 25–29. doi: 10.1007/s11104-010-0638-1.

Caldwell, M. M. and Richards, J. H. (1989) ‘Hydraulic lift: water efflux from upper roots improves effectiveness of water uptake by deep roots’, Oecologia, 79(1), pp. 1–5. doi: 10.1007/BF00378231.

Chen, C. et al. (2004) ‘Row configuration and nitrogen application for Barley–Pea intercropping in Montana’, Agronomy Journal. Madison, WI: American Society of Agronomy, 96, pp. 1730–1738. doi: 10.2134/agronj2004.1730.

Clément, A. et al. (1992) ‘Effects of nitrogen supply and spatial arrangement on the grain yield of a maize/soybean intercrop in a humid subtropical climate’, Canadian Journal of Plant Science. NRC Research Press, 72(1), pp. 57–67. doi: 10.4141/cjps92-007.

Dahmardeh, M. et al. (2009) ‘Effect of intercropping maize (*Zea mays* L.) with cow pea (*Vigna unguiculata* L.) on green forage yield and quality evaluation’, Asian Journal of Plant Sciences, 8(3), pp. 235–239.

Davis, J. H. C. and Woolley, J. N. (1993) ‘Genotypic requirement for intercropping’, Field Crops Research, 34(3), pp. 407–430. doi: https://doi.org/10.1016/0378-4290(93)90124-6.

Dubey, D. N., Kulmi, G. S. and Girish, J. (1995) ‘Relative productivity and economics of sole, mixed and intercropping systems of sorghum (*Sorghum bicolor*) and grain legumes under dryland condition’, Indian Journal of Agricultural Sciences, 65(7), pp. 469–473.

Duchene, O., Vian, J. F. and Celette, F. (2017) ‘Intercropping with legume for agroecological cropping systems: Complementarity and facilitation processes and the importance of soil microorganisms. A review’, Agriculture, Ecosystems and Environment. Elsevier B.V., 240, pp. 148– 161. doi: 10.1016/j.agee.2017.02.019.

Egerton-Warburton, L. M., Querejeta, J. I. and Allen, M. F. (2007) ‘Common mycorrhizal networks provide a potential pathway for the transfer of hydraulically lifted water between plants’, Journal of Experimental Botany, 58(6), pp. 1473–1483. doi: 10.1093/jxb/erm009.

Ghanbari, A. et al. (2010) ‘Effect of maize (*Zea mays* L) - cowpea (*Vigna unguiculata* L) intercropping on light distribution, soil temperature and soil moisture in arid environment’, Journal of Food, Agriculture & Environment, 8(1), pp. 102–108.

Gwathmey, C. O. and Clement, J. D. (2010) ‘Alteration of cotton source-sink relations with plant population density and mepiquat chloride’, Field Crops Research, 116(1–2), pp. 101–107. doi: 10.1016/j.fcr.2009.11.019.

Härdter, R. and Horst, W. J. (1991) ‘Nitrogen and phosphorus use in maize sole cropping and maize/cowpea mixed cropping systems on an Alfisol in the northern Guinea Savanna of Ghana’, Biology and Fertility of Soils, 10(4), pp. 267–275. doi: 10.1007/BF00337377.

Harris, I. et al. (2020) ‘Version 4 of the CRU TS monthly high-resolution gridded multivariate climate dataset’, Scientific Data, 7(1), p. 109. doi: 10.1038/s41597-020-0453-3.

Hauggaard-Nielsen, H. and Jensen, E. S. (2005) ‘Facilitative root interactions in intercrops BT - Root Physiology: from Gene to Function’, in Lambers, H.and Colmer, T. D. (eds). Dordrecht: Springer Netherlands, pp. 237–250. doi: 10.1007/1-4020-4099-7_13.

Hinsinger, P. et al. (2011) ‘P for two, sharing a scarce resource: soil phosphorus acquisition in the rhizosphere of intercropped species’, Plant Physiology, 156(3), pp. 1078–1086. doi: 10.1104/pp.111.175331.

Jakobsen, I., Abbott, L. K. and Robson, A. D. (1992) ‘External hyphae of vesicular-arbuscular mycorrhizal fungi associated with Trifolium subterraneum L.’, New Phytologist. John Wiley & Sons, Ltd (10.1111), 120(3), pp. 371–380. doi: 10.1111/j.1469-8137.1992.tb01077.x.

Jensen, E. S. (1996) ‘Grain yield, symbiotic N2fixation and interspecific competition for inorganic N in pea-barley intercrops’, Plant and Soil, 182(1), pp. 25–38. doi: 10.1007/BF00010992.

Jensen, E. S., Carlsson, G. and Hauggaard-Nielsen, H. (2020) ‘Intercropping of grain legumes and cereals improves the use of soil N resources and reduces the requirement for synthetic fertilizer N: A global-scale analysis’, Agronomy for Sustainable Development, 40(1). doi: 10.1007/s13593-020-0607-x.

Juhos, K., Szabó, S. and Ladányi, M. (2015) ‘Influence of soil properties on crop yield□: a multivariate statistical approach’, International Agrophysics, 29(2), pp. 433–440. doi: 10.1515/intag-2015-0049.

Kaggwa-Asiimwe, R., Andrade-Sanchez, P. and Wang, G. (2013) ‘Plant architecture influences growth and yield response of upland cotton to population density’, Field Crops Research. Elsevier B.V., 145, pp. 52–59. doi: 10.1016/j.fcr.2013.02.005.

Karasawa, T. and Takebe, M. (2012) ‘Temporal or spatial arrangements of cover crops to promote arbuscular mycorrhizal colonization and P uptake of upland crops grown after nonmycorrhizal crops’, Plant and Soil, 353(1–2), pp. 355–366. doi: 10.1007/s11104-011-1036-z.

Lauk, R. and Lauk, E. (2008) ‘Pea-oat intercrops are superior to pea-wheat and pea-barley intercrops’, Acta Agriculturae Scandinavica Section B: Soil and Plant Science, 58(2), pp. 139–144. doi: 10.1080/09064710701412692.

Li, L et al. (2007) ‘Diversity enhances agricultural productivity via rhizosphere phosphorus facilitation on phosphorus-deficient soils’, Proceedings of the National Academy of Sciences of the United States of America, 104(27), pp. 11192–11196. doi: 10.1073/pnas.0704591104.

Li, Long et al. (2007) ‘Diversity enhances agricultural productivity via rhizosphere phosphorus facilitation on phosphorus-deficient soils’, Proceedings of the National Academy of Sciences, 104(27), pp. 3–7. doi: 10.1073/pnas.0704591104.

Li, L. et al. (2014) ‘Plant diversity and overyielding: Insights from belowground facilitation of intercropping in agriculture’, New Phytologist, 203(1), pp. 63–69. doi: 10.1111/nph.12778.

Lithourgidis, A. S. et al. (2007) ‘Sustainable production of barley and wheat by intercropping common vetch’, Agronomy for Sustainable Development, 27(2), pp. 95–99. doi: 10.1051/agro:2006033.

Loreau, M. and Hector, A. (2001) ‘Partitioning selection and complementarity in biodiversity experiments’, Nature, 412(6842), pp. 72–76. doi: 10.1038/35083573.

Ludwig, F. et al. (2004) ‘Below-ground competition between trees and grasses may overwhelm the facilitative effects of hydraulic lift’, Ecology Letters, 7(8), pp. 623–631. doi: 10.1111/j.1461-0248.2004.00615.x.

Mäder, P. et al. (2011) ‘Inoculation of root microorganisms for sustainable wheat-rice and wheat-black gram rotations in India’, Soil Biology and Biochemistry, 43(3), pp. 609–619. doi: 10.1016/j.soilbio.2010.11.031.

Mao, L. et al. (2012) ‘Yield advantage and water saving in maize/pea intercrop’, Field Crops Research. Elsevier B.V., 138, pp. 11–20. doi: 10.1016/j.fcr.2012.09.019.

Martin-Guay, M. O. et al. (2018) ‘The new Green Revolution: Sustainable intensification of agriculture by intercropping’, Science of the Total Environment. Elsevier B.V., 615, pp. 767–772. doi: 10.1016/j.scitotenv.2017.10.024.

Martin, M. P. L. D. and Snaydon, R. W. (1982) ‘Intercropping Barley and Beans I. Effects of Planting Pattern’, Experimental Agriculture. 2008/10/03. Cambridge University Press, 18(2), pp. 139–148. doi: DOI: 10.1017/S0014479700013612.

Mathimaran, N. et al. (2017) ‘Arbuscular mycorrhizal symbiosis and drought tolerance in crop plants’, Mycosphere, 8(3), pp. 361–376. doi: 10.5943/mycosphere/8/3/2.

Mathimaran, N. et al. (2020) ‘Intercropping transplanted pigeon pea with finger millet: Arbuscular mycorrhizal fungi and plant growth promoting rhizobacteria boost yield while reducing fertilizer input’, Frontiers in Sustainable Food Systems, 4(June), pp. 1–12. doi: 10.3389/fsufs.2020.00088.

Moreira, M. Z. et al. (2003) ‘Blackwell Publishing Ltd. Hydraulic lift in a neotropical savanna’, Functional Ecology, 17(1998), pp. 573–581.

Ngwira, A. R., Aune, J. B. and Mkwinda, S. (2012) ‘On-farm evaluation of yield and economic benefit of short term maize legume intercropping systems under conservation agriculture in Malawi’, Field Crops Research. Elsevier B.V., 132, pp. 149–157. doi: 10.1016/j.fcr.2011.12.014.

Oljaca, S. et al. (2000) ‘Effect of plant arrangement pattern and irrigation on efficiency of maize (*Zea mays*) and bean (*Phaseolus vulgaris*) intercropping system’, Journal of Agricultural Science, 135(3), pp. 261–270. doi: 10.1017/S0021859699008321.

Padhi, A. K., Panigrahi, R. K. and Jena, B. K. (2010) ‘Effect of planting geometry and duration of intercrops on performance of pigeonpea-finger millet intercropping systems’, Indian J. Agric. Res., 44(1), pp. 43–47.

Prieto, I. et al. (2011) ‘The role of hydraulic lift on seedling establishment under a nurse plant species in a semi-arid environment’, Perspectives in Plant Ecology, Evolution and Systematics, 13(3), pp. 181–187. doi: 10.1016/j.ppees.2011.05.002.

Querejeta, J. I., Egerton-Warburton, L. M. and Allen, M. F. (2003) ‘Direct nocturnal water transfer from oaks to their mycorrhizal symbionts during severe soil drying’, Oecologia, 134(1), pp. 55–64. doi: 10.1007/s00442-002-1078-2.

Reddy, P. P. (2012) ‘Plant Growth-Promoting Rhizobacteria (PGPR)’, pp. 131–158. doi: 10.1007/978-81-322-0723-8_10.

Saharan, K. et al. (2018) ‘Finger millet growth and nutrient uptake is improved in intercropping With pigeon pea through “Biofertilization” and “Bioirrigation” mediated by arbuscular mycorrhizal fungi and plant growth promoting rhizobacteria’, Frontiers in Environmental Science, 6(June), pp. 1–11. doi: 10.3389/fenvs.2018.00046.

Schipanski, M. E. and Drinkwater, L. E. (2012) ‘Nitrogen fixation in annual and perennial legume-grass mixtures across a fertility gradient’, Plant and Soil, 357(1), pp. 147–159. doi: 10.1007/s11104-012-1137-3.

Schütz, L. et al. (2018) ‘Improving crop yield and nutrient use efficiency via Biofertilization—A global meta-analysis’, Frontiers in Plant Science, 8(January). doi: 10.3389/fpls.2017.02204.

Sekar, J. et al. (2018) ‘Potential of finger millet indigenous rhizobacterium Pseudomonas sp. MSSRFD41 in blast disease management-growth promotion and compatibility with the resident rhizomicrobiome’, Frontiers in Microbiology, 9(MAY), pp. 1–16. doi: 10.3389/fmicb.2018.01029.

Sekiya, N., Araki, H. and Yano, K. (2011) ‘Applying hydraulic lift in an agroecosystem: Forage plants with shoots removed supply water to neighboring vegetable crops’, Plant and Soil, 341(1–2), pp. 39–50. doi: 10.1007/s11104-010-0581-1.

Sekiya, N. and Yano, K. (2002) ‘Water acquisition from rainfall and groundwater by legume crops developing deep rooting systems determined with stable hydrogen isotope compositions of xylem waters’, Field Crops Research, 78(2–3), pp. 133–139. doi: 10.1016/S0378-4290(02)00120-X.

Singh, D. et al. (2019) ‘Bioirrigation: A common mycorrhizal network facilitated the water transfer from deep-rooted pigeon pea to shallow-rooted finger millet under drought’, Plant and Soil, 440, pp. 277–292.

Singh, D. et al. (2020) ‘Deep-rooted pigeon pea promotes the water relations and survival of shallow-rooted finger millet during drought—Despite strong competitive interactions at ambient water availability’, PLoS ONE, 15(2), pp. 1–22. doi: 10.1371/journal.pone.0228993.

Smith, S. E. and Read, D. (2008) ‘17 -Mycorrhizas in agriculture, horticulture and forestry’, in Smith, S.E. and Read, D.B.T.-M.S. (Third E. (eds). London: Academic Press, pp. 611–XVIII. doi: https://doi.org/10.1016/B978-012370526-6.50019-2.

Stenberg, B. (1998) ‘Soil attributes as predictors of crop production under standardized conditions’, Biology and Fertility of Soils, 27(4), pp. 104–112.

Vandermeer, J. H. (1989) The Ecology of Intercropping. Cambridge: Cambridge University Press. doi: DOI: 10.1017/CBO978051162352 .

Willey, R. W. and Osiru, D. S. O. (1972) ‘Studies on mixtures of maize and beans (Phaseolus vulgaris) with particular reference to plant population’, The Journal of Agricultural Science. 2009/03/27. Cambridge University Press, 79(3), pp. 517–529. doi: DOI: 10.1017/S0021859600025909.

Xu, B. C., Li, F. M. and Shan, L. (2008) ‘Switchgrass and milkvetch intercropping under 2:1 row-replacement in semiarid region, northwest China: Aboveground biomass and water use efficiency’, European Journal of Agronomy, 28(3), pp. 485–492. doi: 10.1016/j.eja.2007.11.011.

Zuo, Y. et al. (2004) ‘A study on the improvement iron nutrition of peanut intercropping with maize on nitrogen fixation at early stages of growth of peanut on a calcareous soil’, Soil Science and Plant Nutrition. Taylor & Francis, 50(7), pp. 1071–1078. doi: 10.1080/00380768.2004.10408576.

