## Supplementary material for "Spatial arrangement and biofertilizers enhance the performance of legume – millet intercropping system in rainfed areas of southern India": Supplimentary material

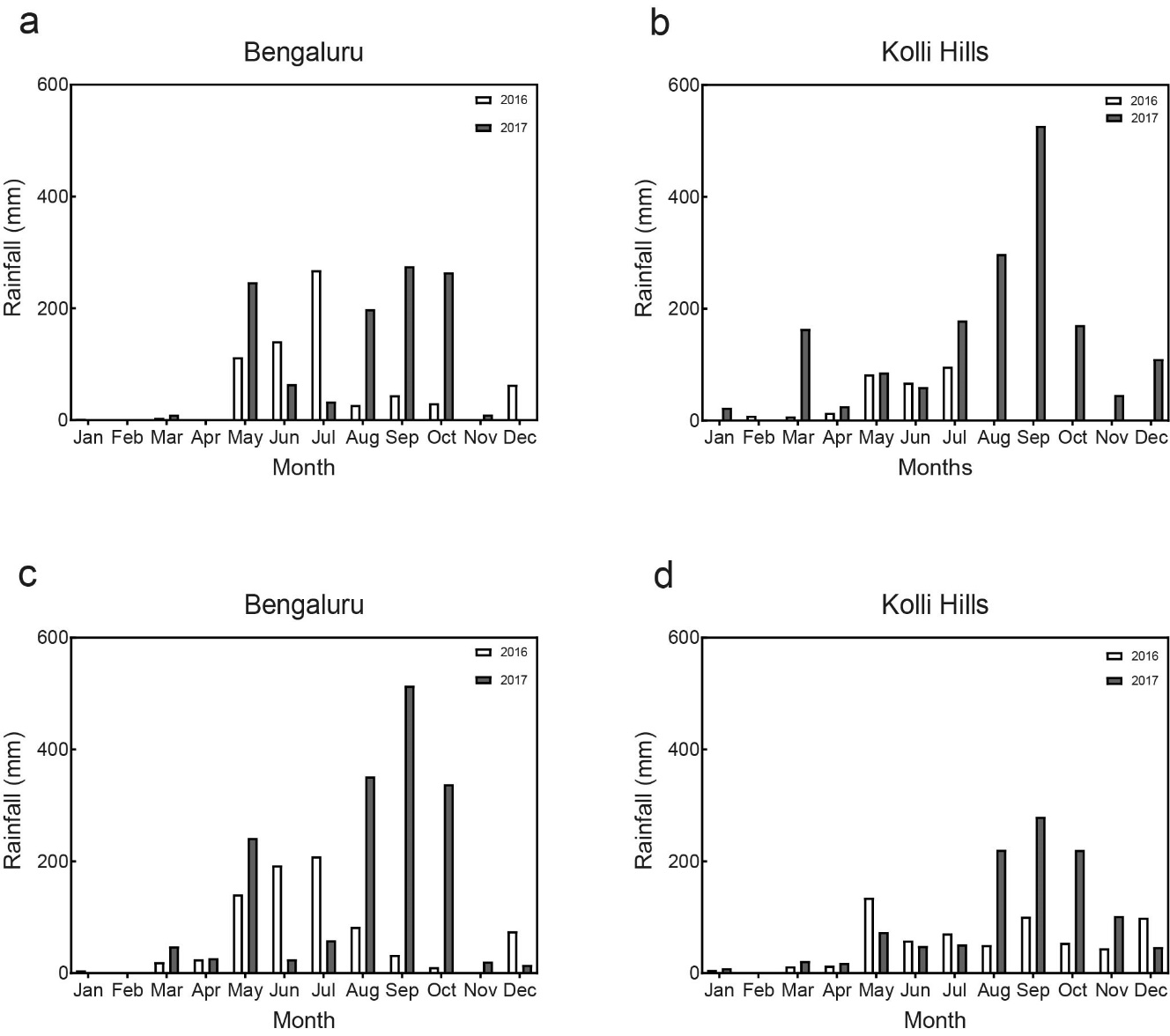

**Fig. S1** Rainfall data of University of Agricultural Sciences, Bengaluru (Figure S1a & S1c) and Kolli Hills, Tamil Nadu India (Figure S1b & S1d) during 2016 and 2017. Data shown in figure S1a and S1b are collected from local weather stations installed at the field site. While, data shown in figure S1c and S1d are observed data from the Climate Research Unit (Harris *et al.*, 2020).

**Table S1** ANOVA table to compare effect of experiment site and treatments, using total biomass, straw and grain biomass from Bengaluru and Kolli Hills field trial from 2016-17. Total biomass includes straw and grain biomass of PP and FM. While, straw biomass represents total straw biomass of PP and FM combined. Similarly, grain biomass represents total grain biomass of PP and FM combined.

| **Total biomass** | **DF** | **SS** | **MS** | **F-value** | **P-value** |
| --- | --- | --- | --- | --- | --- |
| Site | 1 | 5.8590 | 5.85902 | 11.38 | 0.0018 |
| Treatments | 5 | 14.3254 | 2.86508 | 5.57 | 0.0007 |
| Site*Treatment | 5 | 18.7519 | 3.75037 | 7.29 | <0.0001 |
| Error | 36 | 18.5302 | 0.51473 |  |  |
| Total | 47 | 57.4665 |  |  |  |
| **Straw biomass** | **DF** | **SS** | **MS** | **F-value** | **P-value** |
| Site | 1 | 1.7442 | 1.74422 | 10.24 | 0.0029 |
| Treatments | 5 | 4.8492 | 0.96984 | 5.69 | 0.0006 |
| Site*Treatment | 5 | 10.7644 | 2.15288 | 12.64 | <0.0001 |
| Error | 36 | 6.1321 | 0.17034 |  |  |
| Total | 47 | 23.4899 |  |  |  |
| **Grain biomass** | **DF** | **SS** | **MS** | **F-value** | **P-value** |
| Site | 1 | 1.1970 | 1.19701 | 8.51 | 0.0061 |
| Treatments | 5 | 2.6461 | 0.52923 | 3.76 | 0.0077 |
| Site*Treatment | 5 | 1.3798 | 0.27595 | 1.96 | 0.1083 |
| Error | 36 | 5.0664 | 0.14073 |  |  |
| Total | 47 | 10.2893 |  |  |  |

**Table S2** ANOVA table to compare effect of experiment site and treatments, using total biomass, straw and grain biomass from Bengaluru and Kolli Hills field trials data from 2017-18. Total biomass includes straw and grain biomass of PP and FM. While, straw biomass represents total straw biomass of PP and FM combined. Similarly, grain biomass represents total grain biomass of PP and FM combined.

| **Total biomass** | **DF** | **SS** | **MS** | **F-Value** | **P-Value** |
| --- | --- | --- | --- | --- | --- |
| Biofertilization | 1 | 43.471 | 43.4713 | 14.64 | 0.000 |
| Treatments | 4 | 112.438 | 28.1095 | 9.47 | 0.000 |
| Site | 1 | 4.002 | 4.0024 | 1.35 | 0.250 |
| Biofertilization*Treatments | 4 | 6.510 | 1.6274 | 0.55 | 0.701 |
| Biofertilization*Site | 1 | 0.572 | 0.5722 | 0.19 | 0.662 |
| Treatments*Site | 4 | 105.660 | 26.4150 | 8.90 | 0.000 |
| Biofertilization*Treatments*Site | 4 | 12.216 | 3.0541 | 1.03 | 0.400 |
| Error | 60 | 178.172 | 2.9695 |  |  |
| Total | 79 | 463.042 |  |  |  |
| **Straw biomass** | **DF** | **SS** | **MS** | **F-Value** | **P-Value** |
| Biofertilization | 1 | 16.736 | 16.7358 | 6.43 | 0.014 |
| Treatment | 4 | 94.892 | 23.7231 | 9.12 | 0.000 |
| Site | 1 | 19.999 | 19.9991 | 7.69 | 0.007 |
| Biofertilization*Treatment | 4 | 4.349 | 1.0872 | 0.42 | 0.795 |
| Biofertilization*Site | 1 | 0.383 | 0.3826 | 0.15 | 0.703 |
| Treatment*Site | 4 | 90.122 | 22.5304 | 8.66 | 0.000 |
| Biofertilization*Treatment*Site | 4 | 6.498 | 1.6246 | 0.62 | 0.647 |
| Error | 60 | 156.119 | 2.6020 |  |  |
| Total | 79 | 389.098 |  |  |  |
| **Grain biomass** | **DF** | **SS** | **MS** | **F-Value** | **P-Value** |
| Biofertilization | 1 | 6.2617 | 6.2617 | 33.13 | 0.000 |
| Treatment | 4 | 15.4984 | 3.8746 | 20.50 | 0.000 |
| Site | 1 | 6.1080 | 6.1080 | 32.31 | 0.000 |
| Biofertilization*Treatment | 4 | 0.4671 | 0.1168 | 0.62 | 0.652 |
| Biofertilization*Site | 1 | 1.8906 | 1.8906 | 10.00 | 0.002 |
| Treatment*Site | 4 | 5.0820 | 1.2705 | 6.72 | 0.000 |
| Biofertilization*Treatment*Site | 4 | 1.0130 | 0.2533 | 1.34 | 0.266 |
| Error | 60 | 11.3415 | 0.1890 |  |  |
| Total | 79 | 47.6623 |  |  |  |

**Table S3** ANOVA table for total, straw and grain biomass of FM per plant during 2016/17 field trial. Total biomass includes both straw and grain biomass of FM.

| **Total biomass per plant** | **DF** | **SS** | **MS** | **F-Value** | **P-Value** |
| --- | --- | --- | --- | --- | --- |
| Site | 1 | 72.819 | 72.8190 | 47.46 | <0.0001 |
| Treatment | 4 | 65.741 | 16.4353 | 10.71 | <0.0001 |
| Site*Treatment | 4 | 14.598 | 3.6495 | 2.38 | 0.0740 |
| Error | 30 | 46.031 | 1.5344 |  |  |
| Total | 39 | 199.189 |  |  |  |
| **Straw biomass per plant** | **DF** | **SS** | **MS** | **F-Value** | **P-Value** |
| Site | 1 | 36.1190 | 36.1190 | 93.12 | <0.0001 |
| Treatment | 4 | 18.4602 | 4.6151 | 11.90 | <0.0001 |
| Site*Treatment | 4 | 6.7410 | 1.6853 | 4.34 | 0.0069 |
| Error | 30 | 11.6364 | 0.3879 |  |  |
| Total | 39 | 72.9566 |  |  |  |
| **Grain biomass per plant** | **DF** | **SS** | **MS** | **F-Value** | **P-Value** |
| Site | 1 | 6.3680 | 6.36804 | 10.46 | 0.0030 |
| Treatment | 4 | 14.6037 | 3.65094 | 6.00 | 0.0011 |
| Site*Treatment | 4 | 1.8169 | 0.45421 | 0.75 | 0.5685 |
| Error | 30 | 18.2694 | 0.60898 |  |  |
| Total | 39 | 41.0580 |  |  |  |

**Table S4** ANOVA table for total, straw and grain biomass of PP per plant during 2016/17 field trial. Total biomass includes both straw and grain biomass of PP.

| **Total biomass per plant** | **DF** | **SS** | **MS** | **F-Value** | **P-Value** |
| --- | --- | --- | --- | --- | --- |
| Site | 1 | 1019.1 | 1019.09 | 1.43 | 0.2409 |
| Treatment | 4 | 17964.4 | 4491.11 | 6.31 | 0.0008 |
| Site*Treatment | 4 | 9337.3 | 2334.32 | 3.28 | 0.0241 |
| Error | 30 | 21354.1 | 711.80 |  |  |
| Total | 39 | 49674.9 |  |  |  |
| **Straw biomass per plant** | **DF** | **SS** | **MS** | **F-Value** | **P-Value** |
| Site | 1 | 37.5 | 37.54 | 0.14 | 0.7123 |
| Treatment | 4 | 6844.3 | 1711.09 | 6.32 | 0.0008 |
| Site*Treatment | 4 | 3019.8 | 754.96 | 2.79 | 0.0443 |
| Error | 30 | 8128.2 | 270.94 |  |  |
| Total | 39 | 18030.0 |  |  |  |
| **Grain biomass per plant** | **DF** | **SS** | **MS** | **F-Value** | **P-Value** |
| Site | 1 | 665.45 | 665.448 | 5.38 | 0.0274 |
| Treatment | 4 | 2698.58 | 674.646 | 5.46 | 0.0020 |
| Site*Treatment | 4 | 1826.69 | 456.673 | 3.69 | 0.0147 |
| Error | 30 | 3710.10 | 123.670 |  |  |
| Total | 39 | 8900.82 |  |  |  |

**Table S5** ANOVA table for total, straw and grain biomass of FM per plant during 2017-18 field trial. Total biomass includes both straw and grain biomass of FM.

| **Total biomass per plant** | **DF** | **SS** | **MS** | **F-Value** | **P-Value** |
| --- | --- | --- | --- | --- | --- |
| Site | 1 | 1285.67 | 1285.67 | 160.25 | 0.000 |
| Treatment | 3 | 202.91 | 67.64 | 8.43 | 0.000 |
| Biofertilization | 1 | 21.91 | 21.91 | 2.73 | 0.105 |
| Site*Treatment | 3 | 224.50 | 74.83 | 9.33 | 0.000 |
| Site*Biofertilization | 1 | 37.26 | 37.26 | 4.64 | 0.036 |
| Treatment*Biofertilization | 3 | 13.34 | 4.45 | 0.55 | 0.648 |
| Site*Treatment*Biofertilization | 3 | 25.66 | 8.55 | 1.07 | 0.372 |
| Error | 48 | 385.10 | 8.02 |  |  |
| Total | 63 | 2196.35 |  |  |  |
| **Straw biomass per plant** | **DF** | **SS** | **MS** | **F-Value** | **P-Value** |
| Site | 1 | 740.38 | 740.384 | 151.85 | 0.000 |
| Treatment | 3 | 66.13 | 22.044 | 4.52 | 0.007 |
| Biofertilization | 1 | 0.63 | 0.628 | 0.13 | 0.721 |
| Site*Treatment | 3 | 110.34 | 36.779 | 7.54 | 0.000 |
| Site*Biofertilization | 1 | 11.68 | 11.679 | 2.40 | 0.128 |
| Treatment*Biofertilization | 3 | 10.62 | 3.540 | 0.73 | 0.541 |
| Site*Treatment*Biofertilization | 3 | 8.35 | 2.782 | 0.57 | 0.637 |
| Error | 48 | 234.03 | 4.876 |  |  |
| Total | 63 | 1182.16 |  |  |  |
| **Grain biomass per plant** | **DF** | **SS** | **MS** | **F-Value** | **P-Value** |
| Site | 1 | 74.758 | 74.758 | 45.08 | 0.000 |
| Treatment | 3 | 45.212 | 15.071 | 9.09 | 0.000 |
| Biofertilization | 1 | 29.962 | 29.962 | 18.07 | 0.000 |
| Site*Treatment | 3 | 25.016 | 8.339 | 5.03 | 0.004 |
| Site*Biofertilization | 1 | 7.216 | 7.216 | 4.35 | 0.042 |
| Treatment*Biofertilization | 3 | 3.063 | 1.021 | 0.62 | 0.608 |
| Site*Treatment*Biofertilization | 3 | 7.102 | 2.367 | 1.43 | 0.246 |
| Error | 48 | 79.608 | 1.658 |  |  |
| Total | 63 | 271.936 |  |  |  |

**Table S6** ANOVA table for total, straw and grain biomass of PP per plant during 2017/18 field trial. Total biomass includes both straw and grain biomass of PP.

| **Total biomass per plant** | **DF** | **SS** | **MS** | **F-Value** | **P-Value** |
| --- | --- | --- | --- | --- | --- |
| Site | 1 | 343257 | 343257 | 61.85 | 0.000 |
| Treatment | 3 | 119086 | 39695 | 7.15 | 0.000 |
| Biofertilization | 1 | 54798 | 54798 | 9.87 | 0.003 |
| Site*Treatment | 3 | 31971 | 10657 | 1.92 | 0.139 |
| Site*Biofertilization | 1 | 5584 | 5584 | 1.01 | 0.321 |
| Treatment*Biofertilization | 3 | 3371 | 1124 | 0.20 | 0.894 |
| Site*Treatment*Biofertilization | 3 | 8617 | 2872 | 0.52 | 0.672 |
| Error | 48 | 266379 | 5550 |  |  |
| Total | 63 | 833063 |  |  |  |
| **Straw biomass per plant** | **DF** | **SS** | **MS** | **F-Value** | **P-Value** |
| Site | 1 | 353809 | 353809 | 64.35 | 0.000 |
| Treatment | 3 | 122601 | 40867 | 7.43 | 0.000 |
| Biofertilization | 1 | 38317 | 38317 | 6.97 | 0.011 |
| Site*Treatment | 3 | 25252 | 8417 | 1.53 | 0.219 |
| Site*Biofertilization | 1 | 8419 | 8419 | 1.53 | 0.222 |
| Treatment*Biofertilization | 3 | 3302 | 1101 | 0.20 | 0.896 |
| Site*Treatment*Biofertilization | 3 | 6893 | 2298 | 0.42 | 0.741 |
| Error | 48 | 263920 | 5498 |  |  |
| Total | 63 | 822513 |  |  |  |
| **Grain biomass per plant** | **DF** | **SS** | **MS** | **F-Value** | **P-Value** |
| Site | 1 | 79.88 | 79.88 | 2.23 | 0.142 |
| Treatment | 3 | 2059.46 | 686.49 | 19.18 | 0.000 |
| Biofertilization | 1 | 1470.15 | 1470.15 | 41.07 | 0.000 |
| Site*Treatment | 3 | 2105.29 | 701.76 | 19.61 | 0.000 |
| Site*Biofertilization | 1 | 289.85 | 289.85 | 8.10 | 0.006 |
| Treatment*Biofertilization | 3 | 116.45 | 38.82 | 1.08 | 0.365 |
| Site*Treatment*Biofertilization | 3 | 142.42 | 47.47 | 1.33 | 0.277 |
| Error | 48 | 1718.03 | 35.79 |  |  |
| Total | 63 | 7981.52 |  |  |  |

**Table S7** ANOVA table (TWO-WAY ANOVA) of total biomass per plant of FM and PP at Bengaluru, India, in 2017-18 field trial.

| **Total biomass per plant of FM** | **DF** | **SS** | **MS** | **F-Value** | **P-Value** |
| --- | --- | --- | --- | --- | --- |
| Biofertilization | 1 | 45.678 | 45.6780 | 2.81 | 0.1109 |
| Spatial Arrangement | 2 | 145.371 | 72.6854 | 4.47 | 0.0265 |
| Biofertilization*Spatial Arrangement | 2 | 36.050 | 18.0249 | 1.11 | 0.3513 |
| Error | 18 | 292.494 | 16.2497 |  |  |
| Total | 23 | 519.593 |  |  |  |
| **Total biomass per plant of PP** | **DF** | **SS** | **MS** | **F-Value** | **P-Value** |
| Biofertilization | 1 | 8596.1 | 8596.11 | 16.27 | 0.0008 |
| Spatial Arrangement | 2 | 3292.8 | 1646.42 | 3.12 | 0.0688 |
| Biofertilization*Spatial Arrangement | 2 | 2391.0 | 1195.49 | 2.26 | 0.1328 |
| Error | 18 | 9508.2 | 528.23 |  |  |
| Total | 23 | 23788.1 |  |  |  |

**Table S8** ANOVA table (TWO-WAY ANOVA) of total biomass per plant of FM and PP at Kolli Hills, India, in 2017-18 field trial.

| **Total biomass per plant of FM** | **DF** | **Adj SS** | **Adj MS** | **F-Value** | **P-Value** |
| --- | --- | --- | --- | --- | --- |
| Biofertilization | 1 | 0.2993 | 0.2993 | 0.15 | 0.7017 |
| Spatial Arrangement | 2 | 31.6316 | 15.8158 | 8.00 | 0.0033 |
| Biofertilization*Spatial Arrangement | 2 | 3.3286 | 1.6643 | 0.84 | 0.4471 |
| Error | 18 | 35.5713 | 1.9762 |  |  |
| Total | 23 | 70.8307 |  |  |  |
| **Total biomass per plant of PP** | **DF** | **Adj SS** | **Adj MS** | **F-Value** | **P-Value** |
| Biofertilization | 1 | 54247 | 54246.9 | 3.97 | 0.0618 |
| Spatial Arrangement | 2 | 18668 | 9333.8 | 0.68 | 0.5179 |
| Biofertilization*Spatial Arrangement | 2 | 1829 | 914.4 | 0.07 | 0.9355 |
| Error | 18 | 246126 | 13673.7 |  |  |
| Total | 23 | 320869 |  |  |  |

**Table S9** ANOVA table for predawn leaf water potential of FM in 2016-17 and 2017-18 at Bengaluru site. Multifactor ANOVA analysis was performed to find out the effect of different factors and their interaction on water-relations of FM.

| **Predawn leaf water potential (2016/17)** | **DF** | **SS** | **MS** | **F-Value** | **P-Value** |
| --- | --- | --- | --- | --- | --- |
| Treatment | 4 | 3.7650 | 0.941252 | 7.52 | 0.0002 |
| Time | 2 | 1.5043 | 0.752163 | 6.01 | 0.0057 |
| Treatment*Time | 8 | 1.4607 | 0.182585 | 1.46 | 0.2075 |
| Error | 35 | 4.3793 | 0.125123 |  |  |
| Total | 49 | 13.0806 |  |  |  |
| **Predawn leaf water potential (2017/18)** | **DF** | **SS** | **MS** | **F-Value** | **P-Value** |
| Treatment | 3 | 0.31029 | 0.10343 | 7.77 | 0.000 |
| Biofertilizer | 1 | 0.01628 | 0.01628 | 1.22 | 0.274 |
| Time | 2 | 0.41462 | 0.20731 | 15.58 | 0.000 |
| Treatment*Biofertilizer | 3 | 0.21259 | 0.07086 | 5.33 | 0.003 |
| Treatment*Time | 6 | 0.27380 | 0.04563 | 3.43 | 0.007 |
| Biofertilizer*Time | 2 | 0.39849 | 0.19924 | 14.97 | 0.000 |
| Treatment*Biofertilizer*Time | 6 | 0.37823 | 0.06304 | 4.74 | 0.001 |
| Error | 46 | 0.61205 | 0.01331 |  |  |
| Total | 69 | 2.73278 |  |  |  |

**Table S10** Seed weight, plant height, number of panicle per plant, and grain weight per panicle is shown for finger millet from field trial at Bengaluru site during 2016-17.

| Treatments (finger millet) | 1000 seed wt. (g) | Plant height (m) | Panicle/plant | Grain wt./Panicle  (g/panicle) |
| --- | --- | --- | --- | --- |
| T1+ | 3.33 ± 0.19^a^ | 0.71 ± 0.08^a^ | 4.00 ± 0.00^a^ | 13.00 ± 2.31^a^ |
| T3+ | 3.38 ± 0.13^a^ | 0.72 ± 0.03^a^ | 3.75 ± 0.50^a^ | 13.00 ± 1.83^a^ |
| T4+ | 3.35 ± 0.22^a^ | 0.67 ± 0.05^ab^ | 3.25 ± 0.50^ab^ | 8.25 ± 2.22^b^ |
| T5+ | 3.25 ± 0.20^a^ | 0.62 ± 0.05^b^ | 2.75 ± 0.96^b^ | 7.75 ± 4.99^b^ |
| T6+ | 3.26 ± 0.38^a^ | 0.62 ± 0.05^b^ | 3.25 ± 0.50^ab^ | 7.00 ± 1.41^b^ |

**Table S11** Seed weight, plant height, number of pods per plant, and pod weight per plant is shown for pigeon pea from field trial at Bengaluru site during 2016-17.

| Treatments (pigeon pea) | 100 seed wt. (g) | Plant height (m) | Pod/plant | Pod wt./plant (g/plant) |
| --- | --- | --- | --- | --- |
| T2+ | 9.65 ± 0.10^b^ | 1.55 ± 0.01^a^ | 100 ± 7.93^b^ | 108.5 ± 5.32^a^ |
| T3+ | 9.65 ± 0.20^b^ | 1.55 ± 0.12^a^ | 117.75 ± 15.65^a^ | 116.75 ± 6.55^a^ |
| T4+ | 9.70 ± 0.77^b^ | 1.52 ± 0.11^a^ | 112.5 ± 3.70^ab^ | 120.25 ± 32.22^a^ |
| T5+ | 10.27 ± 0.64^ab^ | 1.52 ± 0.05^a^ | 119 ± 1.41^a^ | 119.75 ± 13.15^a^ |
| T6+ | 10.52 ± 0.31^a^ | 1.50 ± 0.05^a^ | 117.5 ± 13.38^a^ | 103 ± 2.16^a^ |

**Table S12** Seed weight, plant height, number of panicle per plant, and grain weight per panicle is shown for finger millet from field trial at Bengaluru during 2017-18.

| Treatments (finger millet) | 1000 seed wt. (g) | Plant height (m) | Panicle/plant | Grain wt./Panicle  (g/panicle) |
| --- | --- | --- | --- | --- |
| T1+ | 2.93 ± 0.15^a^ | 0.98 ± 0.08^a^ | 1 | 3.45 ± 0.42^b^ |
| T1- | 2.90 ± 0.27^a^ | 1.05 ± 0.05^a^ | 1 | 2.97 ± 0.25^b^ |
| T3+ | 3.04 ± 0.11^a^ | 1.04 ± 0.07^a^ | 1 | 5.11 ± 1.29^a^ |
| T3- | 2.83 ± 0.17^a^ | 1.02 ± 0.05^a^ | 1 | 3.44 ± 0.53^b^ |
| T4+ | 3.08 ± 0.10^a^ | 1.02 ± 0.08^a^ | 1 | 3.55 ± 0.35^b^ |
| T4- | 2.93 ± 0.17^a^ | 1.03 ± 0.01^a^ | 1 | 3.28 ± 0.36^b^ |

**Table S13** Seed weight, plant height, number of pods per plant, and pod weight per plant is shown for pigeon pea from field trial at Bengaluru during 2017-18.

| Treatments (pigeon pea) | 100 seed wt. (g) | Plant height (m) | Pod/plant | Pod wt./Plant  (g/plant) |
| --- | --- | --- | --- | --- |
| T1+ | 9.88 ± 0.59^a^ | N/A | N/A | N/A |
| T1- | 9.48 ± 0.67^a^ | N/A | N/A | N/A |
| T3+ | 9.43 ± 0.48^a^ | N/A | N/A | N/A |
| T3- | 9.34 ± 1.14^a^ | N/A | N/A | N/A |
| T4+ | 9.88 ± 0.68^a^ | N/A | N/A | N/A |
| T4- | 9.40 ±0.61^a^ | N/A | N/A | N/A |

**Table S14** Seed weight, plant height, number of panicle per plant, and grain weight per panicle is shown for finger millet from field trial at Kolli Hills during 2016-17.

| Treatments (finger millet) | 1000 seed wt. (g) | Plant height (m) | Panicle/plant | Grain wt./Panicle  (g/panicle) |
| --- | --- | --- | --- | --- |
| T1+ | N/A | 0.74 ± 0.03^a^ | 1.15 ± 0.19^a^ | 8.89 ± 2.63^a^ |
| T3+ | N/A | 0.71 ± 0.11^a^ | 1.58 ± 0.33^a^ | 6.38 ± 0.19^b^ |
| T4+ | N/A | 0.75 ± 0.10^a^ | 1.23 ± 0.26^a^ | 6.02 ± 1.17^b^ |
| T5+ | N/A | 0.70 ± 0.07^a^ | 1.30 ± 0.38^a^ | 6.96 ± 0.46^ab^ |
| T6+ | N/A | 0.69 ± 0.06^a^ | 1.25 ± 0.25^a^ | 7.07 ± 1.75^ab^ |

**Table S15** Seed weight, plant height, number of pods per plant, and pod weight per plant is shown for pigeon pea from field trial at Kolli Hills during 2016-17.

| Treatments (pigeon pea) | 100 seed wt. (g) | Plant height (m) | pod/plant | Pod wt./plant  (g/plant) |
| --- | --- | --- | --- | --- |
| T2+ | 11.80 ± 0.97^a^ | 1.44 ± 0.18^ab^ | 85.50 ± 6.86^b^ | 93.50 ± 20.34^b^ |
| T3+ | 11.10 ± 0.75^a^ | 1.39 ± 0.09^ab^ | 118.75 ± 32.36^ab^ | 112.75 ± 35.75^ab^ |
| T4+ | 12.10 ± 1.44^a^ | 1.46 ± 0.10^a^ | 120.25 ± 24.60^ab^ | 126.00 ± 27.24^ab^ |
| T5+ | 11.40 ± 0.62^a^ | 1.29 ± 0.06^b^ | 158.25 ± 28.93^a^ | 140.00 ± 43.89^a^ |
| T6+ | 10.90 ± 0.97^a^ | 1.30 ± 0.07^b^ | 133.50± 51.80^ab^ | 135.75 ± 14.48^ab^ |

**Table S16** Seed weight, plant height, number of panicle per plant, and grain weight per panicle is shown for finger millet from field trial at Kolli Hills during 2017-18.

| Treatments (finger millet) | 1000 seed wt. (g) | Plant height (m) | Panicle/plant | Grain wt./ear  (g/panicle) |
| --- | --- | --- | --- | --- |
| T1+ | 3.38 ± 0.13^a^ | 1.19 ± 0.08^b^ | 1.35 ± 0.19^ab^ | 3.25 ± 0.67^bc^ |
| T1- | 3.30 ± 0.14^a^ | 0.92 ± 0.03^d^ | 1.05 ± 0.10^d^ | 2.65 ± 0.19^c^ |
| T3+ | 3.40 ± 0.12^a^ | 1.17 ± 0.02^b^ | 1.45 ± 0.10^a^ | 4.32 ± 1.58^a^ |
| T3- | 3.35 ± 0.17^a^ | 1.01 ± 0.04^c^ | 1.25 ± 0.19^abcd^ | 3.09 ± 0.47^bc^ |
| T4+ | 2.45 ± 0.29^b^ | 1.17 ± 0.04^b^ | 1.10 ± 0.12^cd^ | 3.19 ± 0.32^bc^ |
| T4- | 2.68 ± 0.34^b^ | 0.97 ± 0.03^cd^ | 1.30 ± 0.12^abc^ | 2.93 ± 0.31^c^ |
| T5+ | 3.38 ± 0.10^a^ | 1.39 ± 0.02^a^ | 1.20 ± 0.16^bcd^ | 3.04 ± 0.45^bc^ |
| T5- | 3.53 ± 0.21^a^ | 1.15 ± 0.02^b^ | 1.10 ± 0.20^cd^ | 3.96 ± 0.48^ab^ |

**Table S17** Seed weight, plant height, number of pods per plant, and pod weight per plant is shown for pigeon pea from field trial at Kolli Hills during 2017-18.

| Treatments (pigeon pea) | 100 seed wt. (g) | Plant height (m) | Pod/plant | pod wt./plant  (g/panicle) |
| --- | --- | --- | --- | --- |
| T2+ | N/A | 2.75 ± 0.10^abc^ | 147.92 ± 12.76^ab^ | 96.10 ± 10.05^a^ |
| T2- | N/A | 2.70 ± 0.08^abc^ | 134.00 ± 13.45b | 87.88 ± 8.88^a^ |
| T3+ | N/A | 2.81 ± 0.10^a^ | 162.33 ± 19.71^ab^ | 100.93 ± 8.24^a^ |
| T3- | N/A | 2.66 ± 0.05^bc^ | 147.83 ± 43.99^ab^ | 90.75 ± 6.62^a^ |
| T4+ | N/A | 2.76 ± 0.06^ab^ | 151.92 ± 11.74^ab^ | 93.18 ± 7.21^a^ |
| T4- | N/A | 2.75 ± 0.04^abc^ | 132.58 ± 22.93^b^ | 90.82 ± 8.66^a^ |
| T5+ | N/A | 2.64 ± 0.09^c^ | 170.50 ± 19.56^a^ | 100.42 ± 16.23^a^ |
| T5- | N/A | 2.64 ± 0.11^c^ | 138.50 ± 24.89^ab^ | 88.65 ± 12.54^a^ |

Note: Values shown are the average of four replicates ± standard deviation. Tukey`s test (One Way ANOVA) was used for multiple comparison, and values sharing same letters are not significantly different at p>0.05.
